# Large-scale societal dynamics are reflected in human mood and brain

**DOI:** 10.1101/2020.10.07.329201

**Authors:** Alexander V. Lebedev, Christoph Abe, Kasim Acar, Gustavo Deco, Morten L. Kringelbach, Martin Ingvar, Predrag Petrovic

## Abstract

The stock market is a bellwether of socio-economic changes that may directly affect individual well-being. Using large-scale UK-biobank data generated over 14 years, we applied specification curve analysis to rigorously identify significant associations between the local stock market index (FTSE100) and 479,791 UK residents’ mood, as well as their alcohol intake and blood pressure adjusting the results for a large number of potential confounders, including age, sex, linear and non-linear effects of time, research site, other stock market indexes. Furthermore, we found similar associations between FTSE100 and volumetric measures of affective brain regions in a subsample (n=39,755; measurements performed over 5.5 years), which were particularly strong around phase transitions characterized by maximum volatility in the market. The main findings did not depend on applied effect-size estimation criteria (linear methods or mutual information criterion) and were replicated in two independent US-based studies (Parkinson’s Progression Markers Initiative; n=424; performed over 2,5 years and MyConnectome; n=1; 81 measurements over 1,5 years). Our results suggest that phase transitions in the society, indexed by stock market, exhibit close relationships with human mood, health and the affective brain from an individual to population level.

## Introduction

Stock markets mirror the underlying socio-economic status of a population ^1,2^ and may therefore be used as an index or bellwether of the well-being of a society. In line with this idea, previous research has suggested that changes in capital market evolution exhibit a strong impact on traders’ emotional states ^3^, and are associated with the welfare of individuals who have no direct involvement in the stock market ^4^. Moreover, it has been suggested that stock market turbulence is linked to increased anxiety ^5^, self-harm and suicide rates ^6–8^, elevated levels of binge drinking ^9^ and fatal car accidents ^9,10^. These effects may be particularly pronounced in long-lasting financial crises, such as the 2008 stock market crash or the economy slowing in the COVID19 pandemic.

To date, there are no studies that have investigated the association of market behaviour with brain function and structure. In a broader perspective, previous research has suggested that the events that happen in the society have a clear impact on the brain. For example, one study has previously demonstrated how a single extreme aversive global event may impact fear circuits by linking individuals’ geographical proximity to the site of 9/11 terrorist attacks to the reactivation of the amygdala during memory recollection ^11^. Similarly, an upcoming study also suggests that intense experience of the COVID19 outbreak is associated with a volumetric increase of the amygdala ^12^.

The present study aims to understand whether more subtle but frequently occurring global events may leave a trace in the human brain on a population level. Here, we investigated how fluctuations in the stock market are associated with brain structure. Since such fluctuations also mirror global socioeconomic changes in the society ^1^, the investigated associations imply a broader perspective than the specific effects of the market per se.

To do this, we accessed structural MRI data of 39,755 UK citizens from the UK Biobank acquired over approximately 5.5 years (between 2014-05-02 and 2019-10-31), and matched the scan date with the corresponding Financial Times Stock Exchange 100 Index (FTSE100) characterizing stock price of the top 100 UK companies with the largest revenue as our main independent variable (See **Supplement Fig. S1** and **Table S1** for description of the whole dataset, which also included mood data collected over a period of approximately 14 years). The FTSE100 was chosen because the study subjects resided in the UK, and local changes in the economy were expected to impact brain structure on a population level most strongly. In order to index effects on the brain, daily time-series of the market capital index was matched with neuroimaging data focusing on a set of preregistered (https://osf.io/h52gk) brain regions known to play key roles in the processing of rewards and losses, as well as threat and fear ^13–16^: amygdala, nucleus accumbens, insula, anterior, subcallosal and dorsal cingulate and lateral orbitofrontal cortical areas. Abnormal functioning of these circuits has also been documented to play a key role in the pathophysiology of anxiety and depression ^17–20^.

Previous research suggests that brain morphometry is capable of capturing plastic changes that happen after weeks ^21^ or days ^22^ of engagement of the relevant brain networks. Moreover, even acute activation of brain networks is associated with noticeable alterations in morphometric measures ^23^. Even though these changes may represent widely different underlying mechanisms depending on observational time-scales, the literature supports the idea that grey matter changes in major brain networks parallel their functional reorganization ^24^.

Prior to the main analysis, we attempted to replicate previous behavioural findings suggesting a relation of market fluctuations with mood and well-being ^4,7,25,26^ on a large sample from the UK Biobank data (n = 479,791) collected over a period of approximately 14 years. Analysing the relations between FTSE100 and self-reported measures of emotional well-being we confirmed that market ups (higher FTSE100 scores) were associated with higher scores of “happiness” and lower scores in self-reported “negative emotional facets”: irritability, hurt and nervous feelings, anxiety (**Fig. 1; Table 1**). The identified association also held true for the 5.5-years of the MRI subsample (**Supplement Table S2**). We further explored non-imaging variables that are associated with mood changes, i.e. alcohol intake (overall intake frequency and a composite score reflecting weekly intake of all alcoholic beverages) and diastolic blood pressure (automatic readings in mmHg measured at rest), and showed that they were also highly correlated with the FTSE100 (**Fig. 1A**) in that both measures increased when the stock market decreased in value. Several of these effects (relation between stock market and negative emotions, blood pressure or alcohol-intake) were reproduced in the My Connectome data-set consisting of one single subject whose measurements were taken at 81 timepoints during a period or 1,5 years (**Fig. 1B**).

**Fig. 1.**
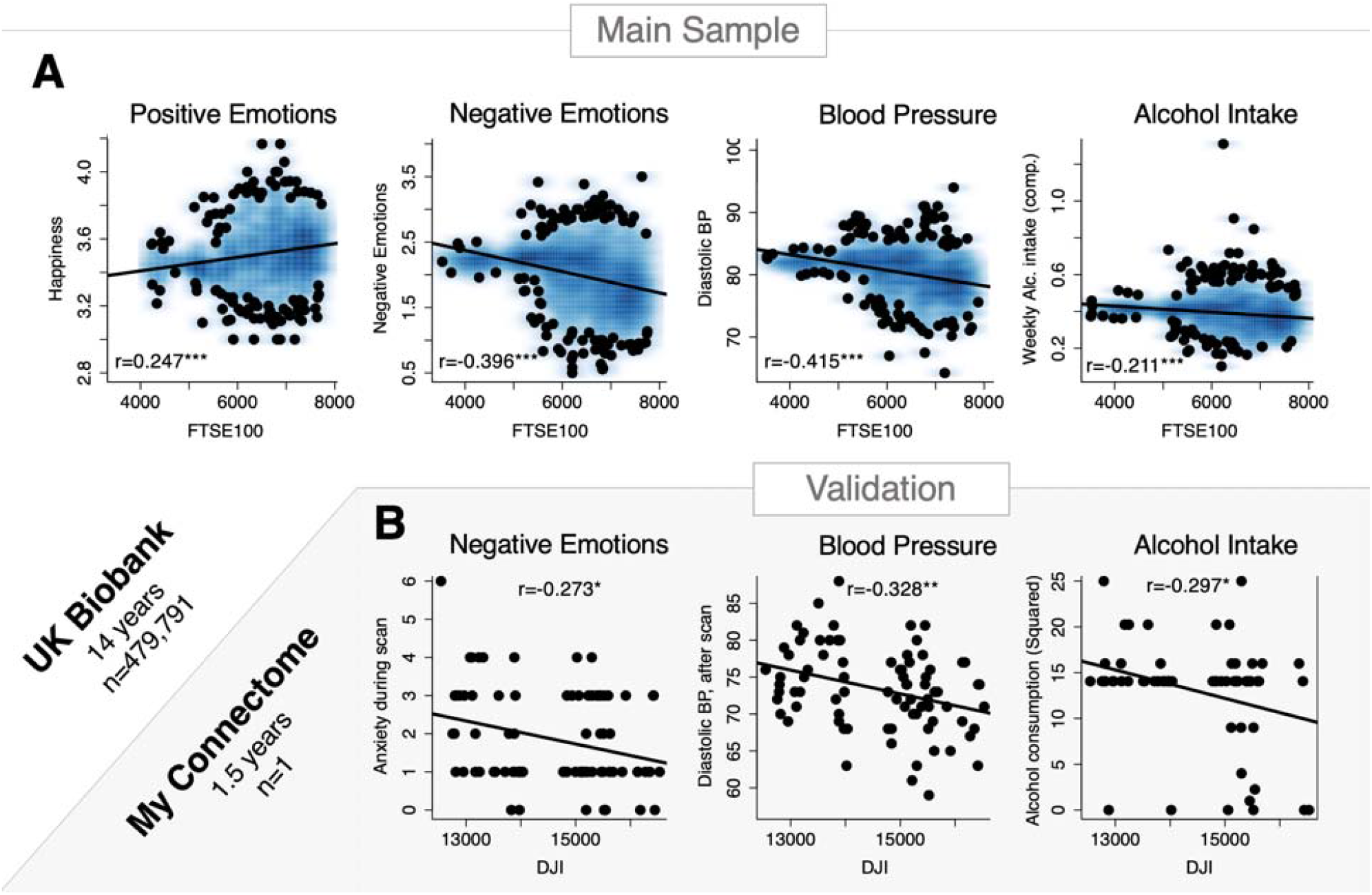
Non-MRI variables and stock market moves. The figure illustrates the identified associations between stock market moves and non-MRI indicators of well-being in the UK Biobank sample (top panel **A**) and My Connectome data, a single-subject study (bottom panel **B**); *p<0.05, **p<0.01, ***p<0.001. Corresponding effect-sizes estimated with mutual information criterion are reported in the supplement (**Supplement Table S11**)

**Table 1.**
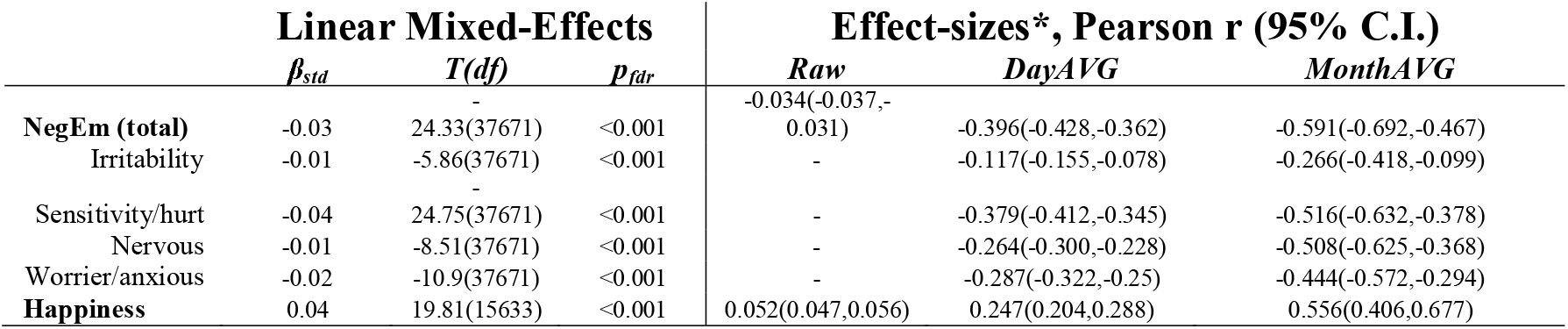
Subjective well-being and FTSE100 scores: 14 years period. β_std_ - standardized β coefficients, p_fdr_ – p-values corrected for multiple testing with false discovery rate. Subcomponents of negative emotions are binary variables (-), Day/MonthAVG – data averaged by days and months. The analyses leveraged random linear mixed effects framework with subject as a random effect, as a subset (n=1427) of the study subjects was assessed twice. * – corresponding effect-sizes estimated with mutual information criterion are reported in the supplement (**Supplement Table S11**).

We then tested and confirmed our main hypothesis by showing that FTSE100 oscillations exhibited significant associations with the morphometry of the affective brain circuits. The most notable result was that bilateral amygdala, involved in threat detection and anxiety processing ^16–20^, showed a negative relation with the UK economic performance (**Fig. 2A**, and **Table 2**, whole-brain analysis revealing that the effects are not limited only by the preregistered regions is reported in **Supplement Fig. S2**). Importantly, this effect was replicated in an independent set of 424 individuals from the PPMI database, an independent clinical study targeting the US population (www.ppmiinfo.org), and conceptually also in “My Connectome” single-subject longitudinal study ^27^. In “My Connectome”, structural data was not publicly available, however, using BOLD-signal variability ^28^ in the amygdala as a proxy biological measure demonstrated that our results also generalise to functional characteristics of the fear network. (**Fig. 2B**). It is worth noting, however, that unlike the main results, detrending the Dow Jones index in these two (PPMI and MyConnectome) datasets reduced effect-sizes without reversing the direction of the associations (**Supplement Fig. S13**).

**Fig. 2.**
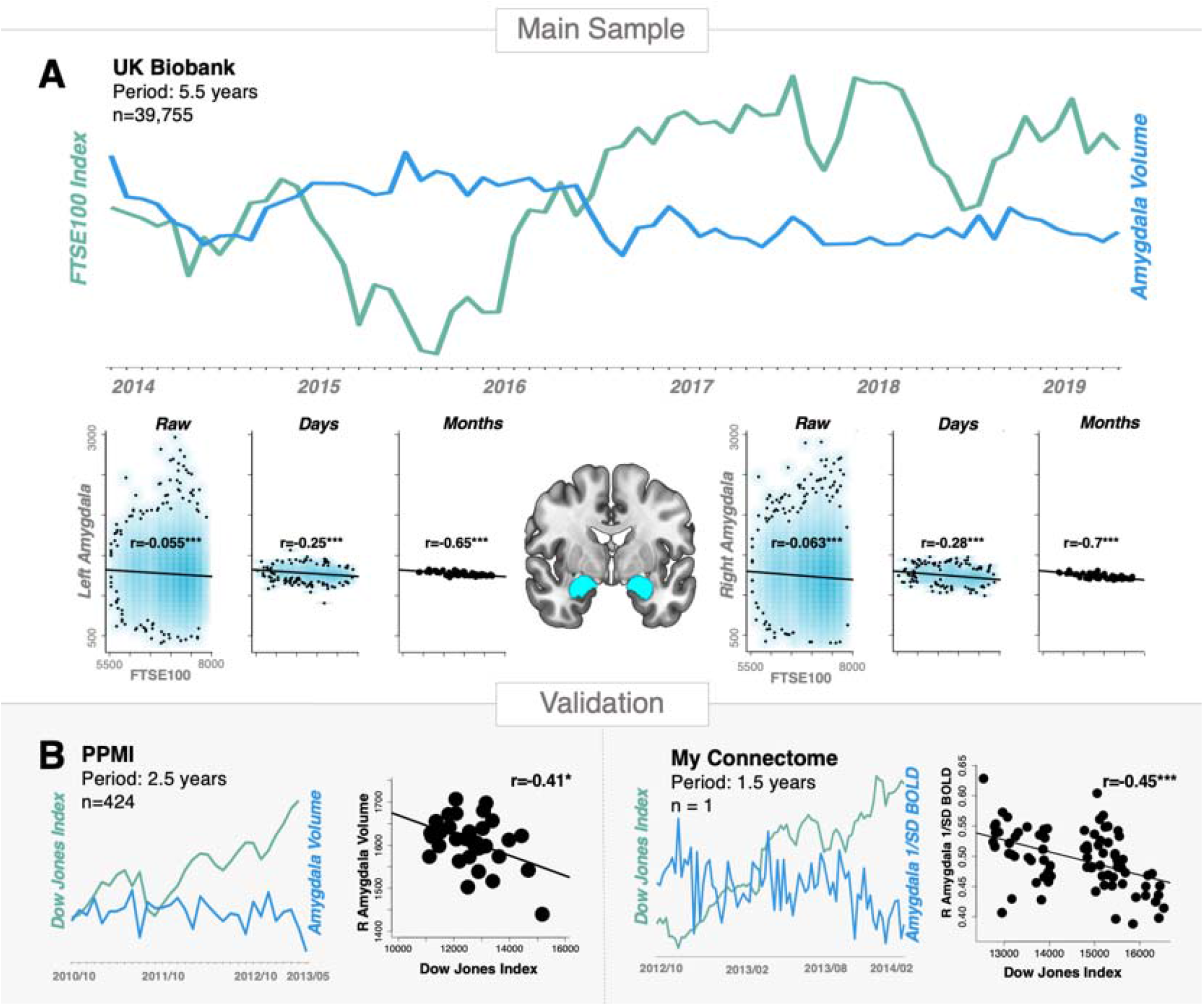
Studied brain-market associations. The figure illustrates the study rationale and reports the investigated effects for the main sample (**A**), as well as their replication (**B**) in a medium-sized (PPMI) and single-subject (My Connectome) fMRI study; *p<0.05, ***p<0.001.

**Table 2.** Associations between FTSE100 and structural characteristics of the fear network: cortical and subcortical volumes. Day/MonthAVG – data averaged over days and months. Intracranial volume (ICV) was selected as a reference measure, which was not expected to exhibit significant associations with global stock market behaviour. β_std_ - standardized β coefficients, p_fdr_ – p-values corrected for multiple testing with false discovery rate. The analyses leveraged random linear mixed effects framework with subject as a random effect, as a subset (n=1427) of the study subjects was scanned twice. * – corresponding effect sizes estimated with the mutual information criterion are reported in the supplement (**Supplement Table S11**)

| Region | Linear Mixed-Effects |  |  | Effect-sizes*, Pearson r (95% C.I.) |  |  |
| --- | --- | --- | --- | --- | --- | --- |
| | $\beta_{std}$ | $T_{883}$ | $p_{idr}$ | Raw, $n=30,775$ | DayAVG, $n=1299$ | MonthAVG, $n=66$ |
| L Amygdala | -0.054 | -9.51 | <0.001 | -0.055(-0.066,-0.043) | -0.253(-0.304,-0.202) | -0.615(-0.746,-0.439) |
| R Amygdala | -0.062 | -10.91 | <0.001 | -0.063(-0.074,-0.052) | -0.282(-0.332,-0.231) | -0.644(-0.767,-0.477) |
| L Accumbens | -0.054 | -9.54 | <0.001 | -0.055(-0.066,-0.044) | -0.232(-0.283,-0.18) | -0.623(-0.752,-0.449) |
| R Accumbens | -0.062 | -10.89 | <0.001 | -0.064(-0.075,-0.052) | -0.259(-0.309,-0.207) | -0.662(-0.779,-0.5) |
| L LOFC | -0.026 | -4.68 | <0.001 | -0.031(-0.042,-0.02) | -0.141(-0.193,-0.087) | -0.443(-0.619,-0.225) |
| R LOFC | -0.019 | -3.49 | 0.001 | -0.023(-0.034,-0.012) | -0.082(-0.136,-0.028) | -0.292(-0.499,-0.054) |
| L Insula | 0.037 | 6.62 | <0.001 | 0.042(0.031,0.053) | 0.21(0.157,0.261) | 0.494(0.286,0.657) |
| R Insula | 0.032 | 5.86 | <0.001 | 0.037(0.026,0.048) | 0.187(0.134,0.239) | 0.413(0.19,0.595) |
| **L Subcallosal** | 0.018 | 3.18 | 0.002 | 0.02(0.009,0.032) | 0.134(0.08,0.187) | 0.322(0.087,0.524) |
| **R Subcallosal** | 0.015 | 2.74 | 0.008 | 0.019(0.007,0.03) | 0.129(0.075,0.182) | 0.305(0.068,0.509) |
| **L Anterior Cingulate** | 0.025 | 4.67 | <0.001 | 0.035(0.024,0.046) | 0.178(0.124,0.23) | 0.408(0.184,0.591) |
| **R Anterior Cingulate** | 0.024 | 4.44 | <0.001 | 0.033(0.022,0.044) | 0.169(0.116,0.222) | 0.428(0.207,0.607) |
| **L Paracingulate** | 0.004 | 0.8 | 0.426 | 0.001(-0.01,0.013) | 0.044(-0.01,0.098) | 0.106(-0.14,0.339) |
| **R Paracingulate** | 0.005 | 0.83 | 0.426 | 0.003(-0.008,0.014) | 0.052(-0.003,0.106) | 0.122(-0.124,0.353) |
| **Intracranial Volume** | 0.004 | 0.87 | 0.426 | -0.009(-0.02,0.002) | -0.016(-0.07,0.038) | -0.098(-0.332,0.147) |

Splitting the study timeline into 6 equal periods (11 months each) we showed that the correlations are strongest during and following phase transition events, i.e. when the change and variability of stock market dynamics is most pronounced (**Supplement Fig S8**).

Similar findings were observed for nucleus accumbens and lateral orbitofrontal cortex (lOFC) that also increased in volume when market decreased (**Fig. 3, A** and **B**). While nucleus accumbens is mostly known for being involved in reward anticipation, it is equally important for processing losses ^13,14^. lOFC has been suggested to be involved in processing expectations within the emotional domain ^29–31^, including losses and rewards ^32,33^. Further supporting this, a significant interaction (β = -0.01, t_776_ = -2.87, p=0.004, p_fdr_ = 0.05) between FTSE100 and income index was found on the right lOFC volume (**Supplement Table S5**). Post-hoc analyses revealed the highest effects in individuals with the lowest and highest income, suggesting that right lOFC of those subjects is particularly sensitive to the capital market swings. Insula and anterior cingulate showed the opposite effect, i.e. the size correlated positively with the market (**Fig. 3, A** and **C**).

**Fig. 3.**
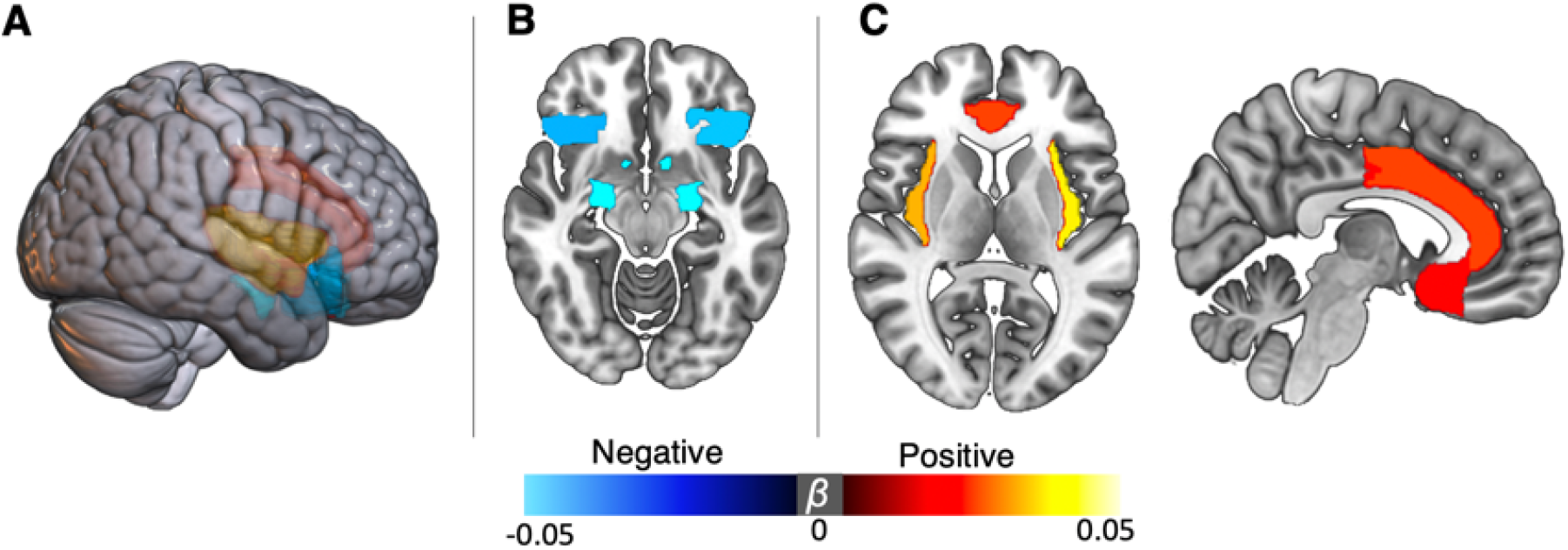
Regional profile of brain-market associations. **A**) Three-dimensional view of the significant findings (p_FDR_<0.05). FTSE100 exhibited negative. **B**) associations with amygdala, nucleus accumbens and orbitofrontal cortex, whereas insular and cingulate regions were positively. **C**) associated with the index scores. The analyses leveraged random linear mixed effects framework with subject as a random effect, as a subset (n=1427) of the study subjects was scanned twice.

All regions mentioned above, except for lOFC, are involved in both positive and negative emotional processes ^13–15^. Out of those regions, subcortical nuclei (amygdala and nucleus accumbens) correlated negatively with the stock market. In contrast, the cortical regions (insula and anterior cingulate) seem exhibit a positive relation with the stock market moves. The magnitude of the identified effects varied depending on time scale with median Pearson correlation |*r*| = 0.033 (0.001-0.064) for the raw data, |*r*| = 0.169 (0.017-0.282) for the day-averaged measures, and |*r*| = 0.492 (0.09-0.73) when brain and market data were averaged over months (**Table 2**). Importantly, all of the reported associations changed very little after detrending the FTSE100 time-series. Deconvolving FTSE100 time-series into low- and high-frequency domains using fast Fourier transform, showed that low-frequency oscillations mostly drive the effect, although, a similar pattern of associations was observed for the high frequency band (**Supplement Fig. S3, Table S6**).

We amended the preregistered protocol by adding additional possible confounding variables to confirm that the main results are robust and withstand correction for age, sex, presence of psychiatric diagnoses, seasonal effects (months) and intracranial volume (**Supplement Table S3**). Moreover, robustness of our findings was also confirmed in the specification curve analysis ^34^ that showed stability of the effects with respect to different model specification strategies (**Supplement Figure S10, S11**).

When considering the indexes of the UK’s fifteen top trading partners ^35^, a similar pattern of associations as the one for FTSE100 was observed for the equivalent local European indexes (e.g. German GDAXI, Dutch AEX, French FCHI) but was of smaller magnitude (**Fig. 4**). The associations further declined or had different directions for markets that were more distant in a socioeconomical dimension (as also reflected in a weaker correlation with FTSE100), including the reference Shanghai Composite Index (SSEC). Importantly, the results also withstood correction for these indexes (**Supplement Table S4**), which implies that the local economic performance captured by the FTSE100 exhibits a specific association with the characteristics of the scanned UK population.

**Fig. 4.**
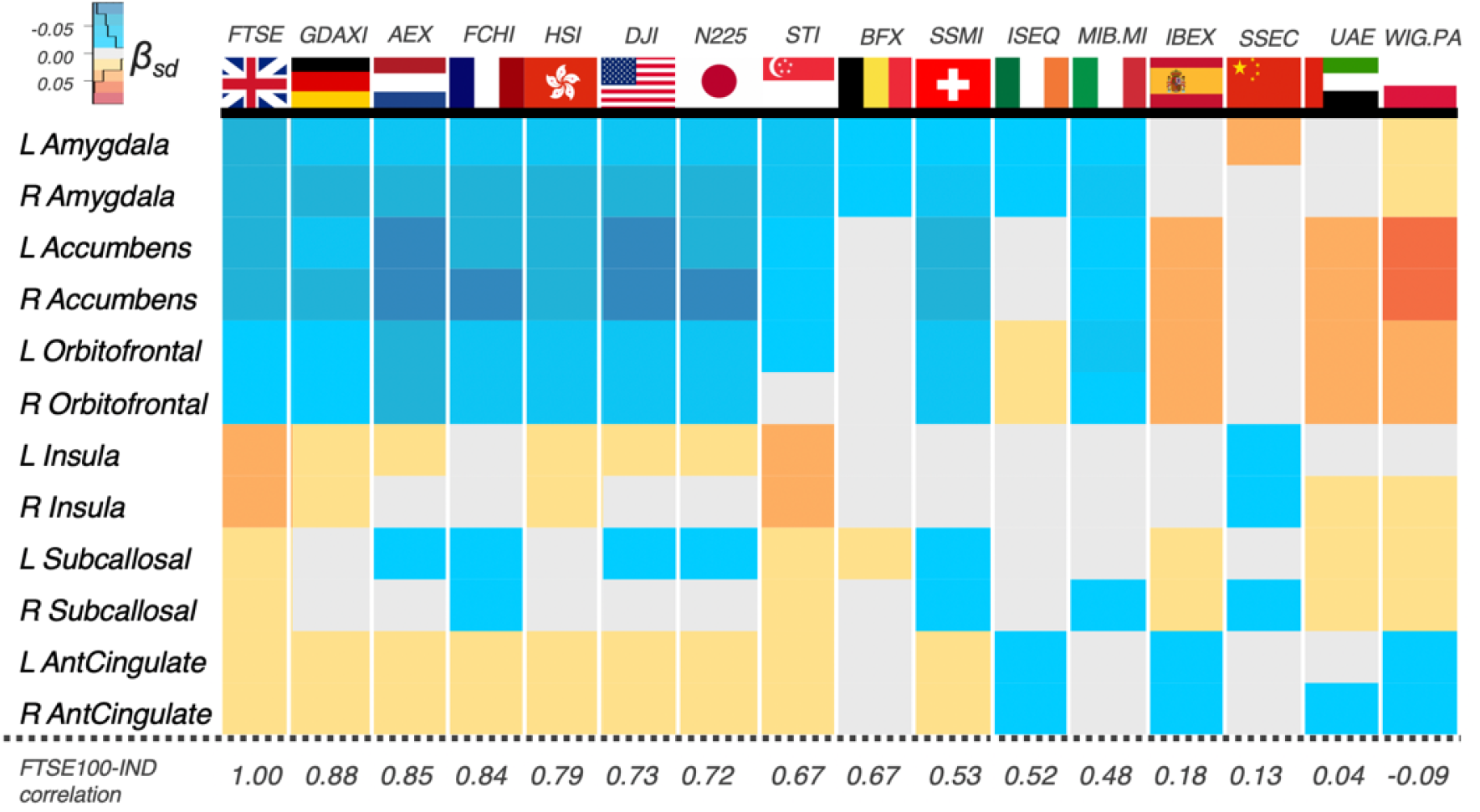
Pattern of brain-market associations for different capital market indexes. Strongest associations were found for the UK market index (FTSE100). Japanese and Singapore and Hong Kong indexes also exhibited a similar pattern of associations possibly reflecting socioeconomic and geographic similarity with the UK, whereas Dow Jones Industrial Average (DJA) likely reflects major contribution of the United States to the world economy. Chinese index (preregistered as a reference) had one of the weakest associations with the studied volumetric measures. FTSE100-IND correlations: Pearson correlation of FTSE100 with other investigated indexes. The analyses leveraged random linear mixed effects framework with subject as a random effect, as a subset (n=1427) of the study subjects was scanned twice.

Regarding causality, the most widely accepted hypothesis states that population mood and well-being are impacted by market via effects on the socioeconomic environment ^4,6,26^. These effects, heavily reinforced by media, represent an index for variables like fluctuations in house prices ^36^ and unemployment ^37^. They can also be considered as threat signals that subsequently impact brains and emotional states of the population ^10^. Another related hypothesis from socionomics is currently growing in popularity. It puts forward the idea of “social mood” as a herding-driven emergent state that originates from population dynamics and subsequently drives global processes, including economic crises, wars, art and fashion ^1,2^. According to this hypothesis, social mood is an inherently hidden state of the society. It is related (but not identical) to the mood of individuals that such a group consists of. This hypothesis is conceptually supported by the data acquired in small-scale experimental studies demonstrating involvement of reward and fear circuits in future financial decisions ^38–40^. Of importance for the present discussion, this hypothesis considers stock market dynamics as a valuable “metric stick” of the social mood and global societal dynamics ^1^.

To begin to further investigate these relationships, we evaluated associations with time-lagged Pearson correlation. We identified that brain volumes correlate higher with *earlier* market prices. The correlation remains significant for approximately one year and then gradually decays (**Fig. 5**). While an autocorrelation, as expected, is present in the stock market time-series ^41^ (**Supplement Table S7**), the fact that earlier economic data peaks with the brain volume implies that the market events may be antecedent to the brain volume fluctuations, offering initial evidence that the market “impacts” the brain, mood, and well-being. The same analyses were carried out on the monthly scale yielding similar results (**Supplement Fig. S4**) and also for the mood data with the FTSE100, although no clear antecedent relationship could be drawn for the latter (**Supplement Fig. S5**).

**Fig. 5.**
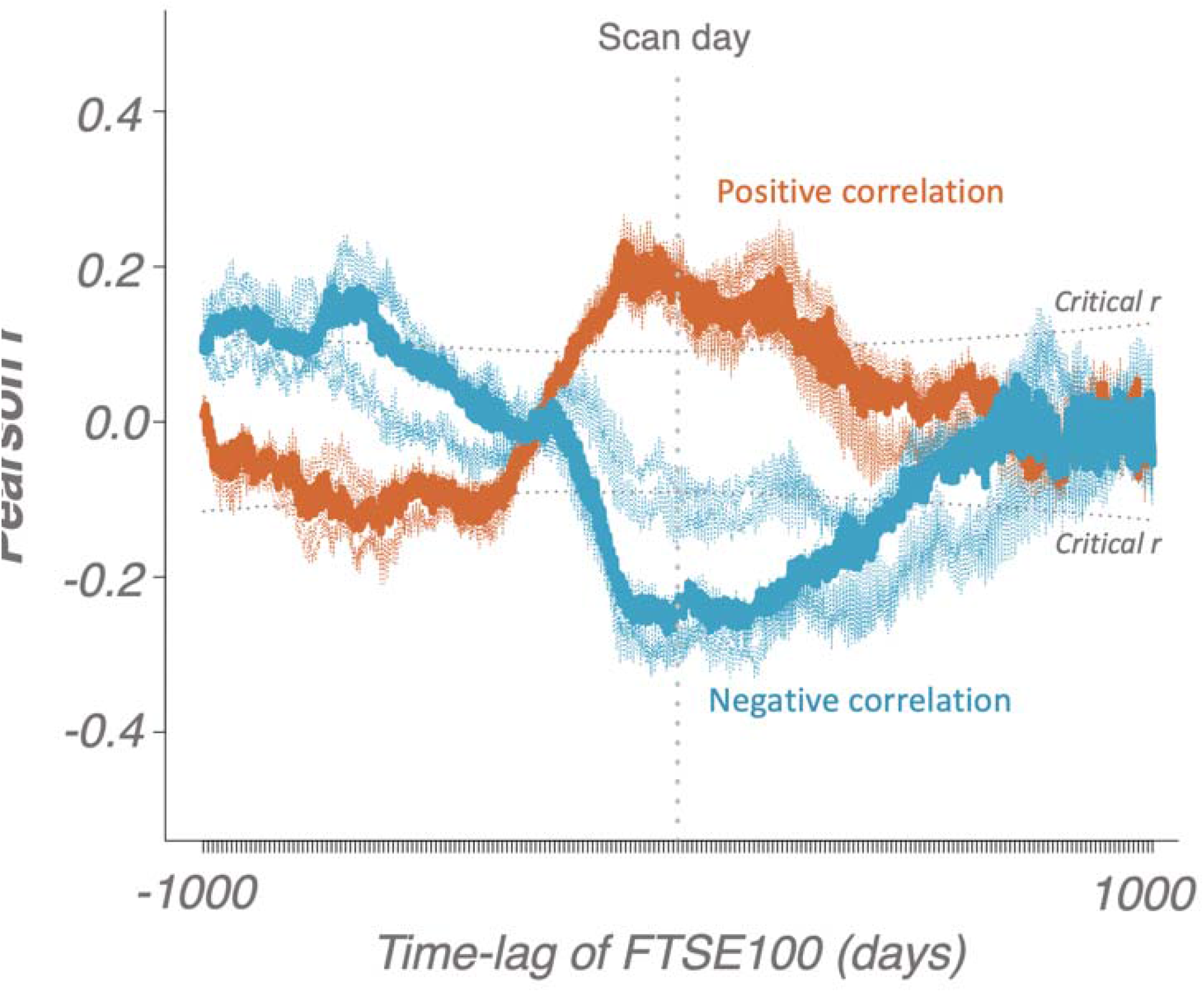
Pearson correlations for the brain and FTSE100-lagged data averaged over days. Transparent lines represent individual regions whereas thick lines represent medians of the correlations. Dotted boundaries represent critical r-values for α=0.001. The plot represents magnitudes of associations between brain data at the date of scanning and the FTSE100 index shifter forward (right) and backward (left) in time. Note a reversed peak for earlier dates reflective of autocorrelations.

Despite the fact that Toda-Yamamoto implementation of Granger Causality tests specifically designed for serially correlated data ^42^ provided somewhat stronger support in favour of a causal link “Market impacts Population Brain/Mood” (**Supplement Tables S8 and S9**), a caution is still advised when interpreting these analyses due to scale-free properties of the investigated time-series (**Fig 6**). To illustrate this point, we first show the absence of any significant effects after shuffling the dates (**Supplement Table S10**, column 1), but appearance of residual associations for the time-shifted data (**Supplement Table S10**, column 2, also seen on **Fig 5**). Importantly, simulating stock market data with 1/f noise is capable of producing effect-sizes of similar magnitude (**Supplement Table S10**, column 3), pooled effect of which, however, converges to zero due to inconsistency of directions in the estimated associations (***Supplement Fig. S12***), and, unlike the main results, also disappear after adjusting for other stock market indexes (**Supplement Table S10**, column 4), confirming that the main effect is not driven by a randomly-seeded 1/f noise. Moreover, we demonstrated that the magnitude of the brain-market links (measured as median squared root correlations) is related to economic and sociocultural ties of the UK to other countries ^35,43^ (**Supplement Fig S6 and S7**) and that no other global candidate metrics with 1/f properties (UK seismic activity and mortality rates) exhibit an equivalent level of specificity with respect to the investigated variables (**Supplement Fig S9**).

**Fig 6.**
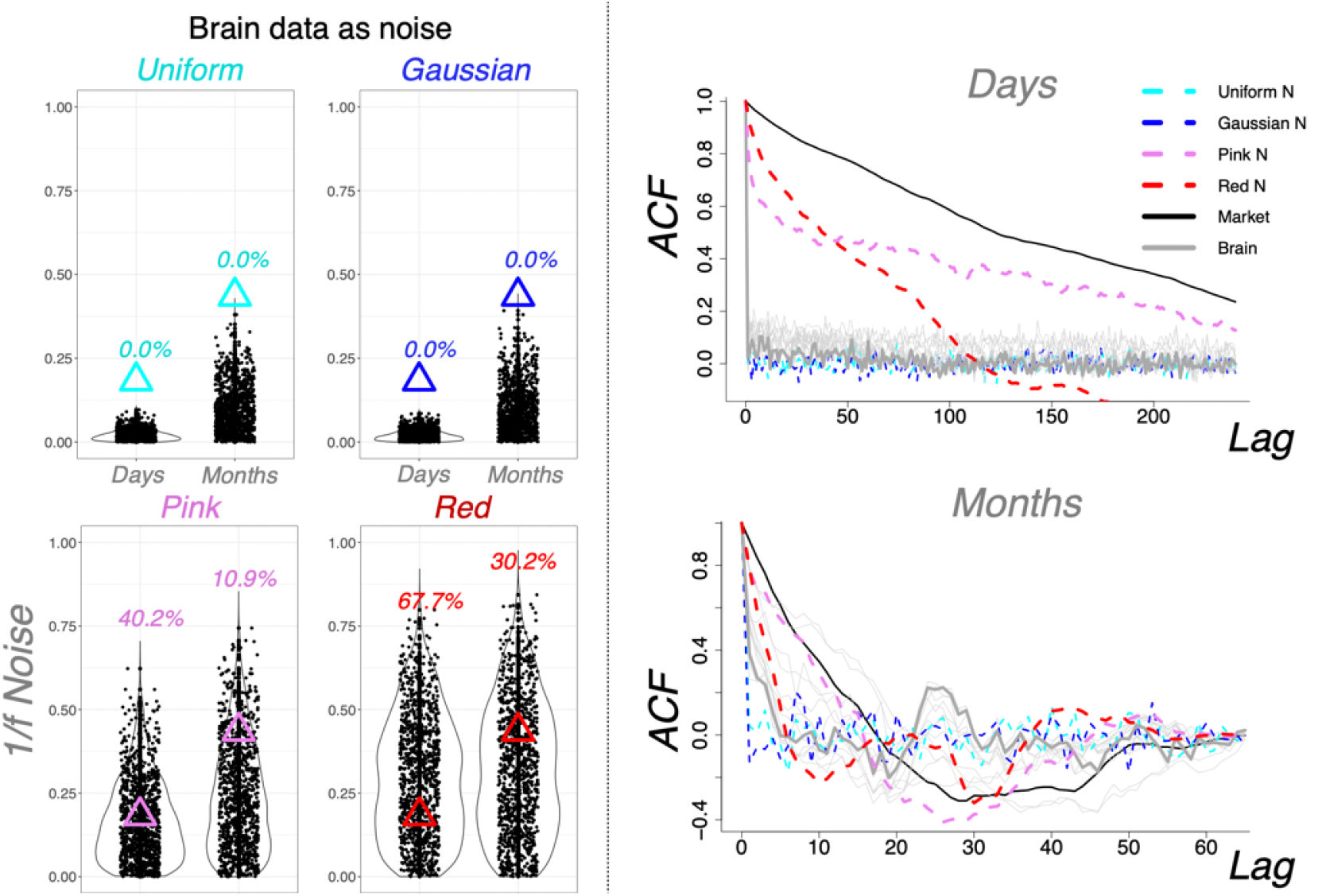
Noise simulation experiments and autocorrelation function density plots. LEFT: Uniform and gaussian noise simulations failed to produce the effect sizes of equivalent (rootsquared) magnitude to the one found in the present study (top). However, 1/f noise was capable of inducing such associations (bottom). Note that we intentionally used root-squared estimates to illustrate these effects. Without this step, all of the estimates from multiple noise simulations converge to zero (**Supplement Fig. S12**), unlike the reported results showing consistent directionality in different time-bins and three independent samples. RIGHT: Autocorrelation function (ACF) density plots demonstrating scale-free properties of the stock market data most similar to the ones of 1/f noise (pink and red).

Due to self-similarity properties identified in the data (**Fig 6**, right panel), we decided to conduct a follow-up series of noise simulation experiments. Simulating brain data with uniform and gaussian noise failed to induce the afore-mentioned correlations with FTSE100, but, as expected, they were more likely to be discovered for the brain data simulated with 1/f noise (**Fig 6**, left panel).

Therefore, it appears so that scale-free properties are observed at different levels of population dynamics, which is reflected in fluctuations of stock markets, mood and brains. To confirm that the effects still hold after accounting for scale-free noise, we repeated the simulations of brain data with 1/f noise and matched it with the stock markets of the UK’s 15 top trading partners. We then subtracted the yielded Pearson correlations from the real ones (prior to calculating the medians) and, as expected, the effect sizes only became larger (**Supplement Fig S7 B**). Moreover, a negative association was also identified for a number of sociocultural distances of the UK from 17 countries using data from Liu et al, 2018 ^43^ (**Supplement Fig S7**). All of the above supported Casti’s hypothesis of stock markets as a useful metric stick for global societal dynamics ^1^.

There is a number of important considerations that must be taken into account when interpreting our results. First, violation of random sampling assumptions can potentially occur in large-scale datasets collected over long time-periods, including the UK Biobank data ^44^. It is worth noting, however, that our results survived all of the undertaken adjustments for potential analytical biases and potential confounds, including research site, age, sex, linear and non-linear effects of time, patient status, other stock market indexes, intracranial volume, as well as all possible combinations of the selected confounds in the specification curve analyses ^34^. However, if these assumptions are, indeed, consistently violated across different time-bins and dataset scales, resulting in the same effects in different samples, including longitudinal single-subject studies, this is already an important finding implying that the investigated effects are big enough to impact complex behavioural patterns, including enrolment likelihood of individuals with certain psychological and biological traits, or at the very least represent an important confound that must be taken into account when designing any studies (cross-sectional or longitudinal) that use data collected over long periods ^27^. The same caution applies to potential presence of scanner drifts. And whilst it is theoretically possible for them to exhibit similar dynamics to the one of stock markets, we address this limitation by showing similar associations with the non-MRI (behavioural) data.

Another important point of discussion is the topic of randomness and the origin of scale-free noise in complex systems. Modern studies suggest that the stock market behaviour should not be modelled as a ‘random walk’ (i.e. having Gaussian distribution), but rather as a non-Gaussian process with random ‘jumps’ resulting in fat-tailed distributions ^45^. In such cases, leveraging linear methods to estimate associations between two variables may not represent the best solution. Recognizing importance of this point we conducted a set of analyses for varying time-windows and also replicated our results using mutual information criterion, which, as some may argue, may be a more potent strategy for detecting these non-linearities.

Finally, we would like to highlight again that the investigated market variable (FTSE100 index) does not represent the stock market per se, but rather reflects a current socioeconomic state of the society.

Our study presented evidences for self-organized criticality present in stock market behaviour supporting the socionomic hypothesis of “social mood” as a driving factor in global societal processes. Here we show that these dynamics may originate in scale-free temporal dynamics present in many complex systems in nature ^41^.

Despite being small on an individual level, these effects may have a large influence on a population level, as the previous studies have suggested ^4–8,10^. This is underscored by our objective measure of diastolic blood pressure that in average differed five units between the samples measured during the lowest market outcomes compared with the ones measured during the highest market outcomes. This effect may have clinical relevance on a population level. Moreover, our results suggest that some sub-populations are particularly vulnerable to economic turbulences, such as individuals with low and very high income. Understanding these complex but nevertheless important processes is of crucial relevance for sustainable and well-being-oriented economic development 46,47.

## Supplementary Materials

### Materials and Methods

#### Preregistration

The study was conducted in accordance with the Declaration of Helsinki on Ethical Principles for Medical Research Involving Human Subjects ^48^ and after approval of the submitted research proposal by the UK Biobank (Application ID #62895) was preregistered at the Open Science Foundation Framework database (https://osf.io/h52gk) prior to data transfer. The primary submitted hypothesis was that “global market fluctuations exhibit significant associations with structure and function of brain regions of the fear circuit”. The present set of analyses was fully focused on the structural data.

#### Subjects

The current project targeted a population of British citizens from the UK Biobank. The main sample consisted of 41,182 data-points from a total of 39,755 UK citizens who completed MRI sessions at least once and was assessed over a period of approximately 4.5 years (between 2014-05-02 and 2019-10-31). The larger sample consisted of 547,005 data-points (479,791 individuals) and was used in the analyses of mood-market relationships (See **Table S1** for descriptive statistics of the two samples).

#### MRI data

Brain scans were collected on a 3 Tesla scanner Siemens Magnetom Skyra Syngo MR D13. Structural T1 3D scans were collected adhering to a standardized protocol as described in the UK Biobank materials (see https://www.fmrib.ox.ac.uk/ukbiobank/protocol/index.html): TA: 4:54, voxel size:1.0×1.0×1.0 mm.

Structural scans were preprocessed employing automated steps as implemented in FSL, yielding region-specific measures of cortical and subcortical volumes parcellated according to the Harvard-Oxford atlas (https://fsl.fmrib.ox.ac.uk/fsl/fslwiki/Atlases).

The main analysis was focused on 14 regions-of-interest that were selected in advance as the ones playing major roles in affective processing: amygdala, nucleus accumbens, insula, anterior, subcallosal and dorsal cingulate and lateral orbitofrontal cortical areas.

#### Market Data

Historical daily time-series data of FTSE100 stock market index was extracted from yahoo finance website (https://finance.yahoo.com/) and matched with fluctuations of structural brain measures of the studied population on each of the scanning days.

The preregistered market outcome was The Financial Times Stock Exchange 100 Index (FTSE100), a widely accepted metric of UK economic performance characterizing stock price of the top 100 UK companies with biggest revenue.

For the analyses we used the daily adjusted close value that represents the closing price after adjustments for all applicable splits and dividend distributions adhering to Center for Research in Security Prices (CRSP) calculation standards. No additional transformations were applied on the extracted time-series in the results reported in the main text. However, the stability of the investigated associations was later confirmed on the de-trended time-series. As a part of the exploratory analyses, we also looked into low- and high-frequency bands of the market time-series deconvolved with fast Fourier transform.

#### Analysis

Initially, we planned to employ classical regression methods to estimate linear relationships between brain and stock market data, followed by a correction for potential confounders. However, after discovering that we will receive access to subjects that were scanned twice (n=1427) we decided to take advantage of this by employing methods of linear mixed-effects modelling introducing “subject” as a random effect variable. All of the main analytical steps, however, were further repeated with classical regression methods confirming the presented results.

Causality tests were applied on the primary outcomes to investigate two alternative causal paths of the studied processes: H1: “brain impacts market” and H2: “market impacts brain”. The workflow adhered to the Toda-Yamamoto implementation of Granger Causality for non-stationary data ^42^ and consisted of: 1) testing for integration and determining max order of integration, 2) setting up a VAR-model in levels for the non-differenced data, 3) determining the lag length, 4) Portmanteau test for residual serial correlation, 5) adding the maximum order of integration to the number of lags, generating the augmented VAR-model, 6) application of the Wald χ^2^ test of the two alternative augmented models: “market impacts brain” and “brain impacts market”.

All statistical analyses were performed using R programming language. Linear mixed-effects models and Toda-Yamamoto Granger causality analyses were implemented using ‘nlme’ and ‘vars’ packages, respectively.

**Fig. S1.**
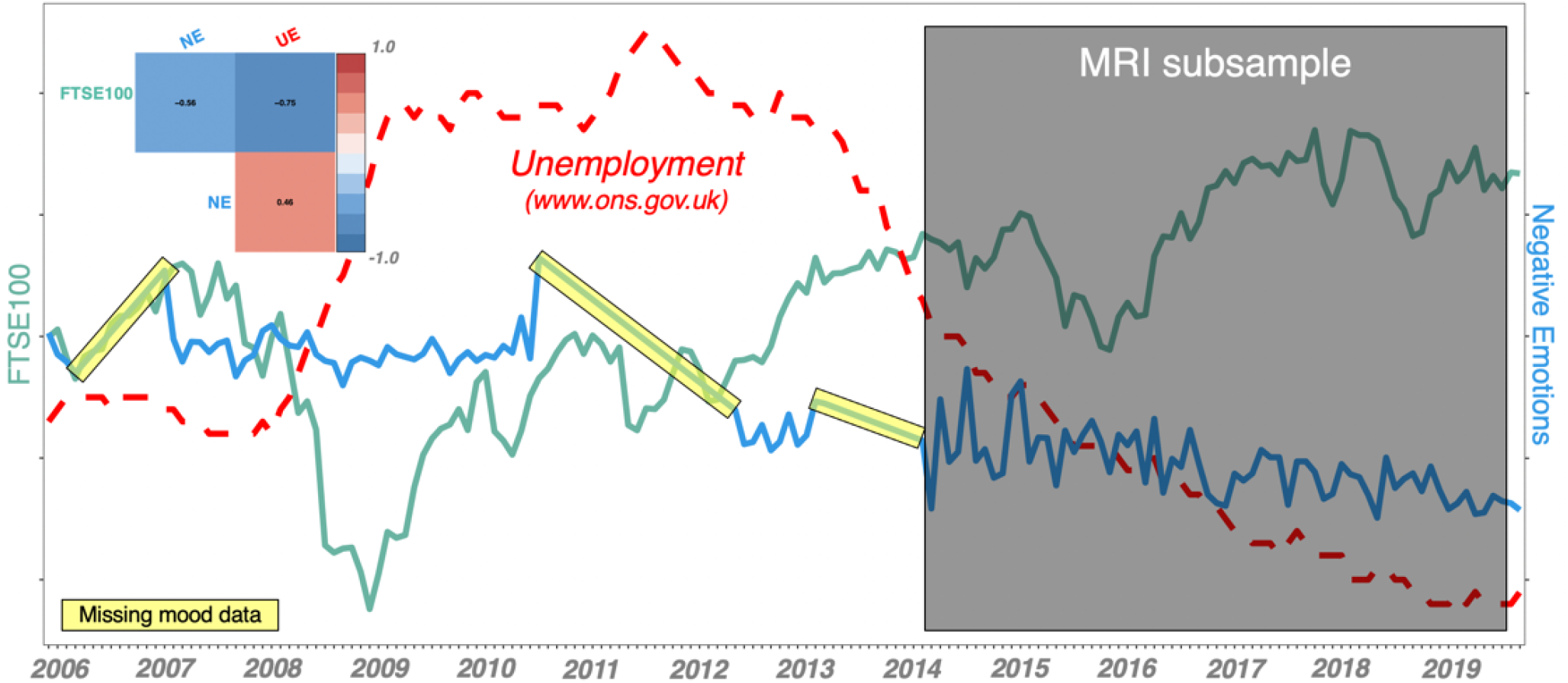
Illustration of the investigated period. Mood data was available for a larger period of approximately 14 years and was missing for a number of time points (interpolated on the figure, but not in the main analyses). The plot illustrates dynamics of the UK stock market (FTSE100), unemployment levels (UE, source: http://www.ons.gov.uk and self-reported negative emotions (NE) for this period. MRI data was available for a period of approximately 5.5 years.

**Fig. S2.**
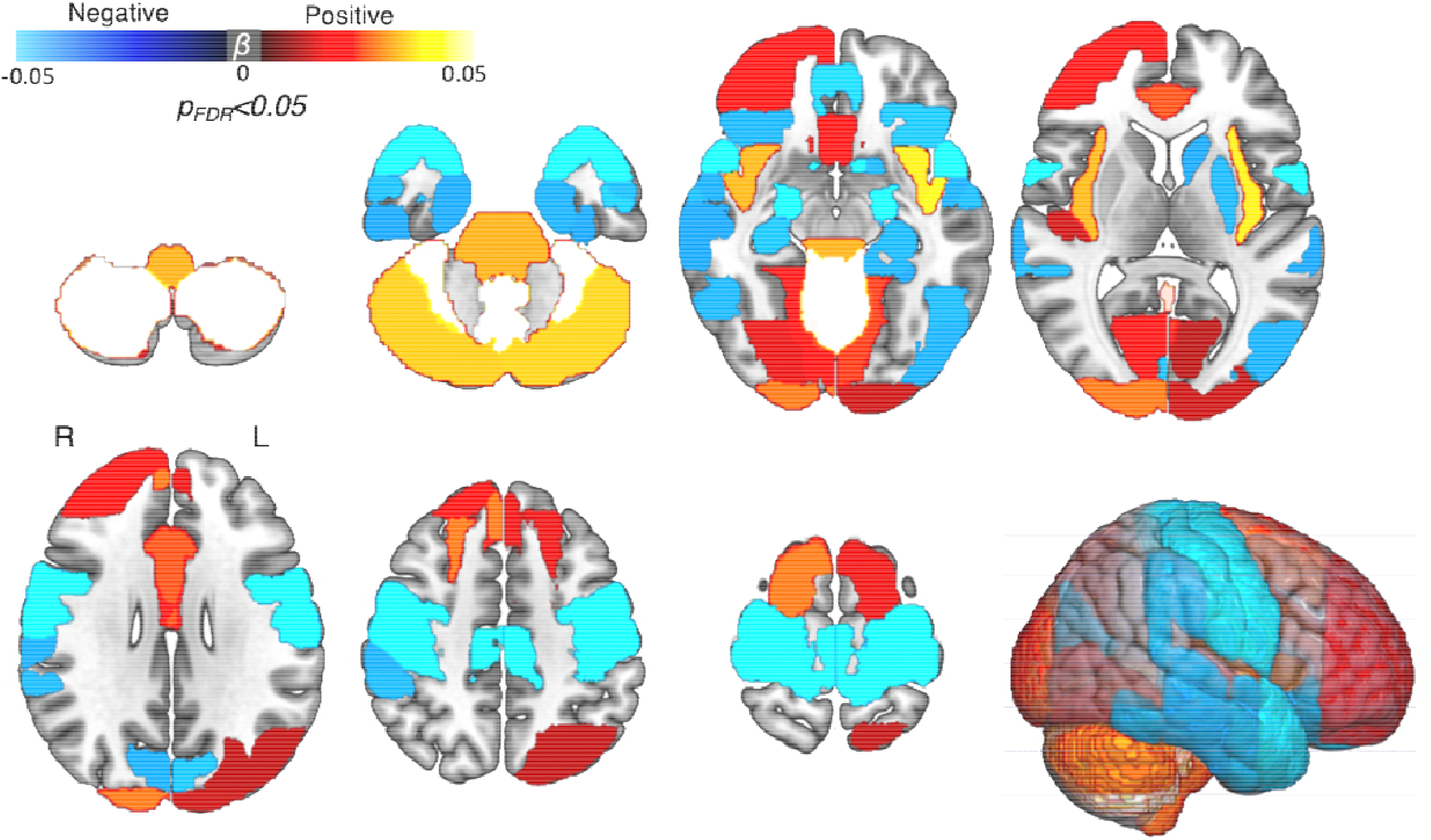
Associations between FTSE100 and grey matter volumes: whole-brain analysis. The associations were estimated employing mass-univariate strategy following correction for multiple testing with false discovery rate. Whilst the effects were not specific to the selected regions that are proven components of the fear/reward network, it can be noted that the largest effects are, indeed, seen in the areas playing key roles in affective and motor processing.

**Fig. S3.**
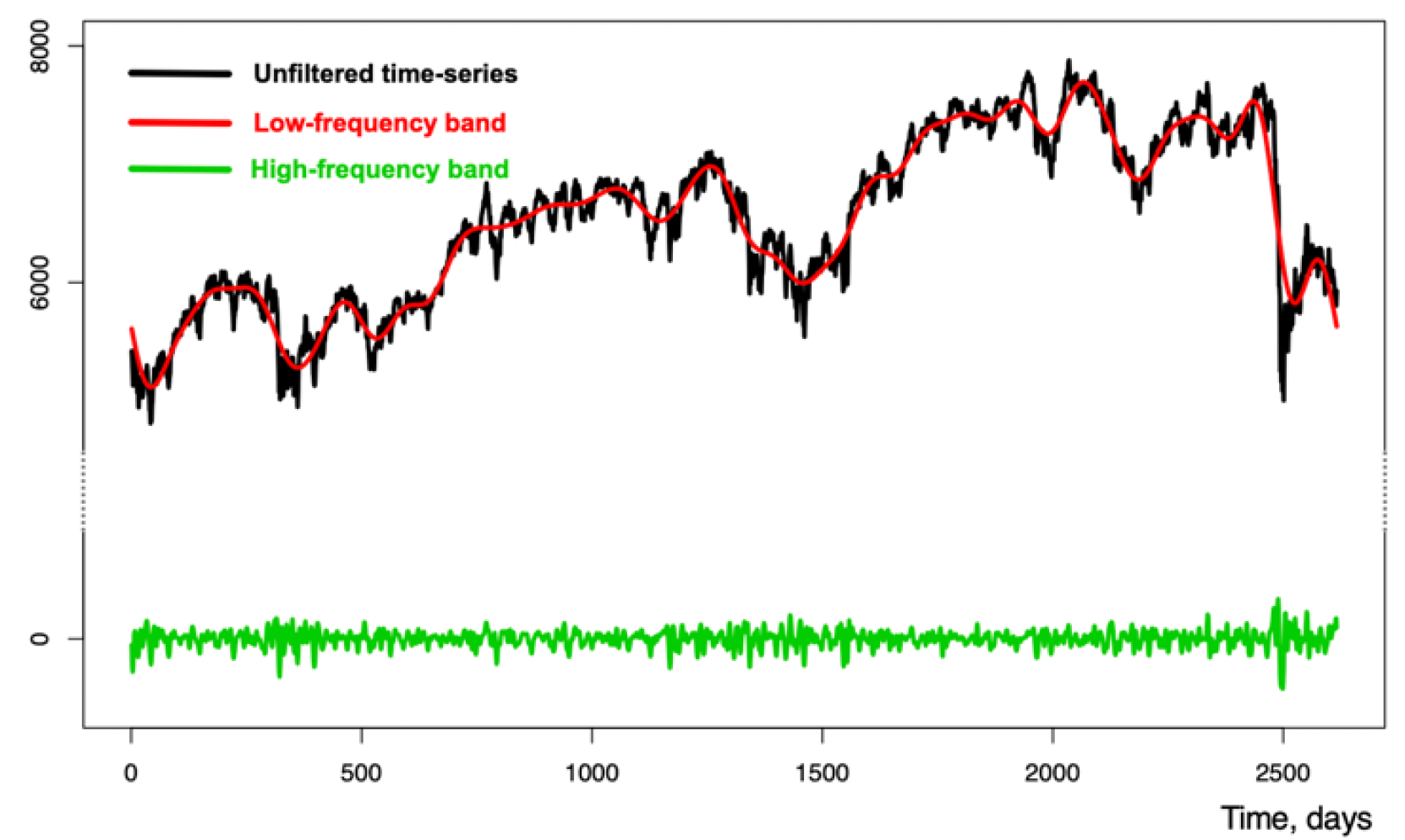
Frequency band analysis. FTSE100 time-series deconvolved with Fast Fourier Transform (FFT) into low- and high-frequency bands, which were further analyzed in relation to the brain and mood data employing methods of linear modeling.

**Figure S4.**
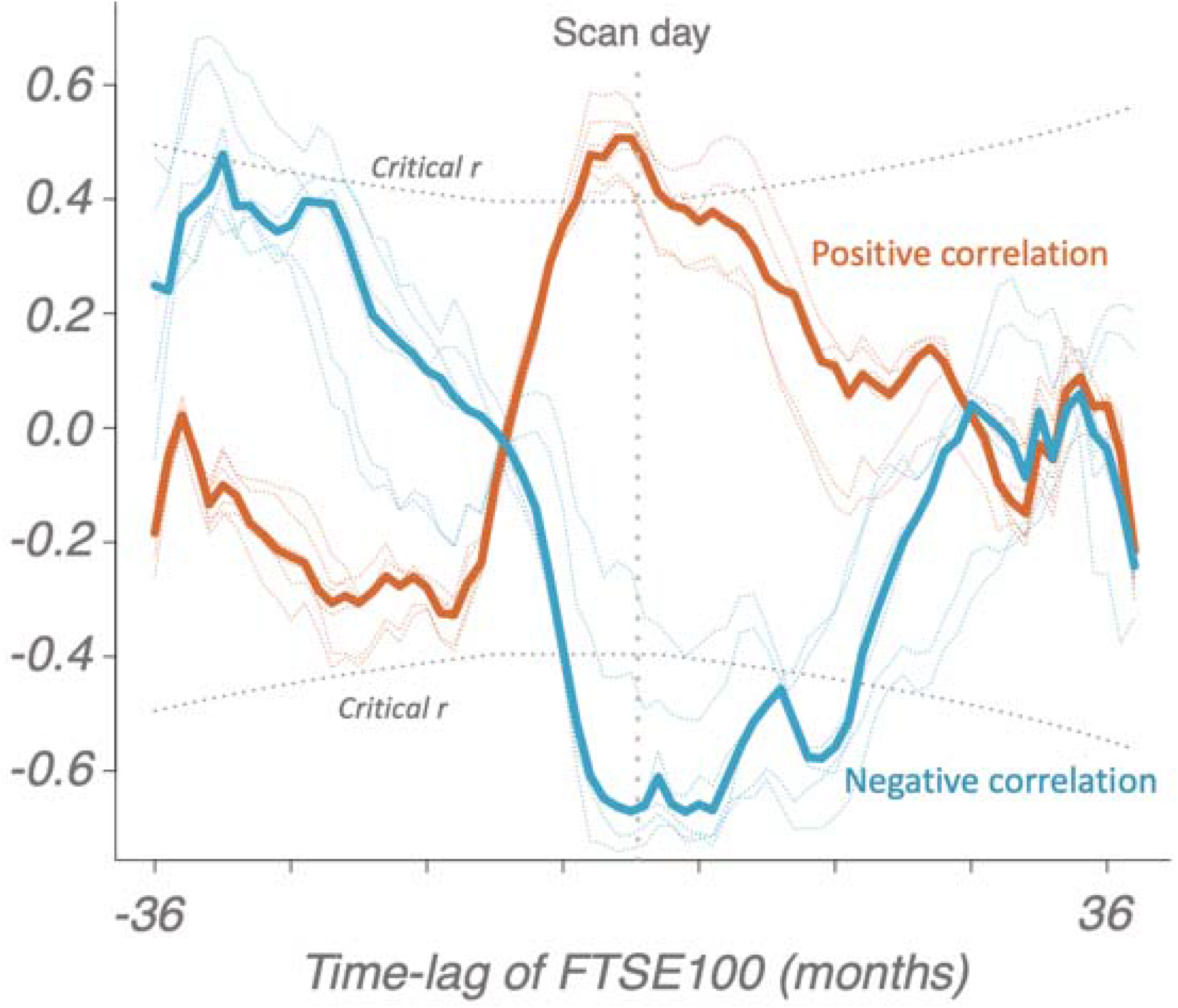
Pearson correlations for the brain and FTSE100-lagged data averaged over months. Transparent lines represent individual regions whereas thick lines represent medians of the correlations. Dotted boundaries represent critical r-values for α=0.001.

**Figure S5.**
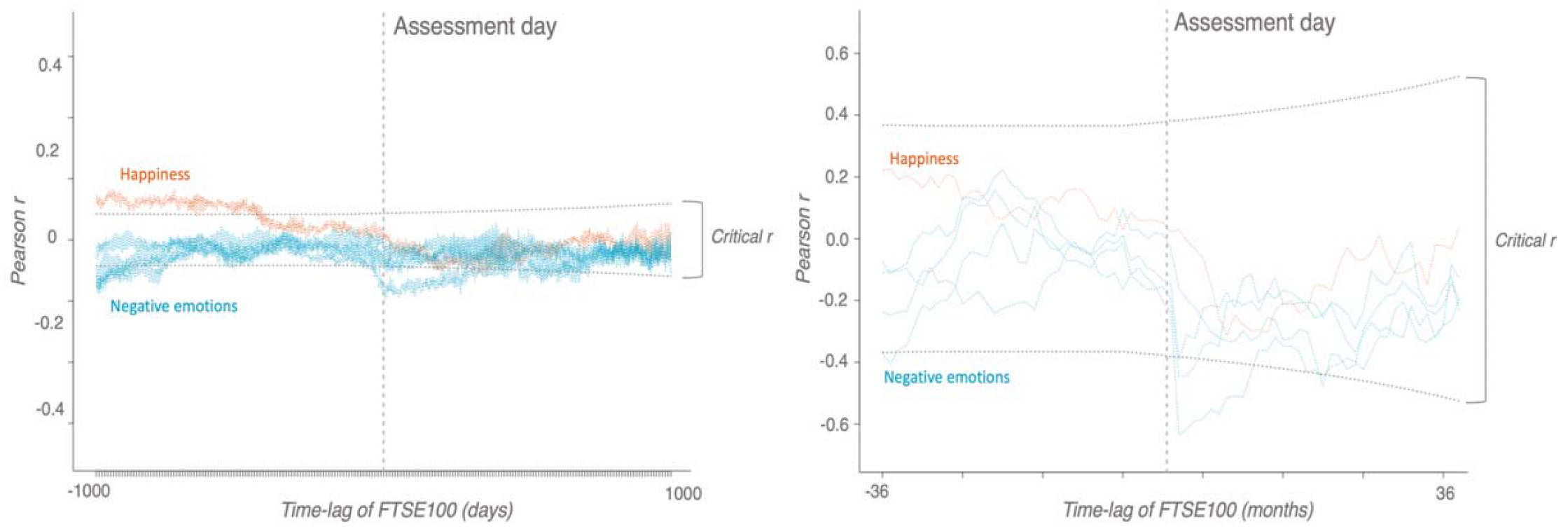
Correlations over days and months. Pearson correlations for the mood and FTSE100-lagged data averaged over days (left) and months (right). Dotted boundaries represent critical r-values for α=0.001.

**Figure S6.**
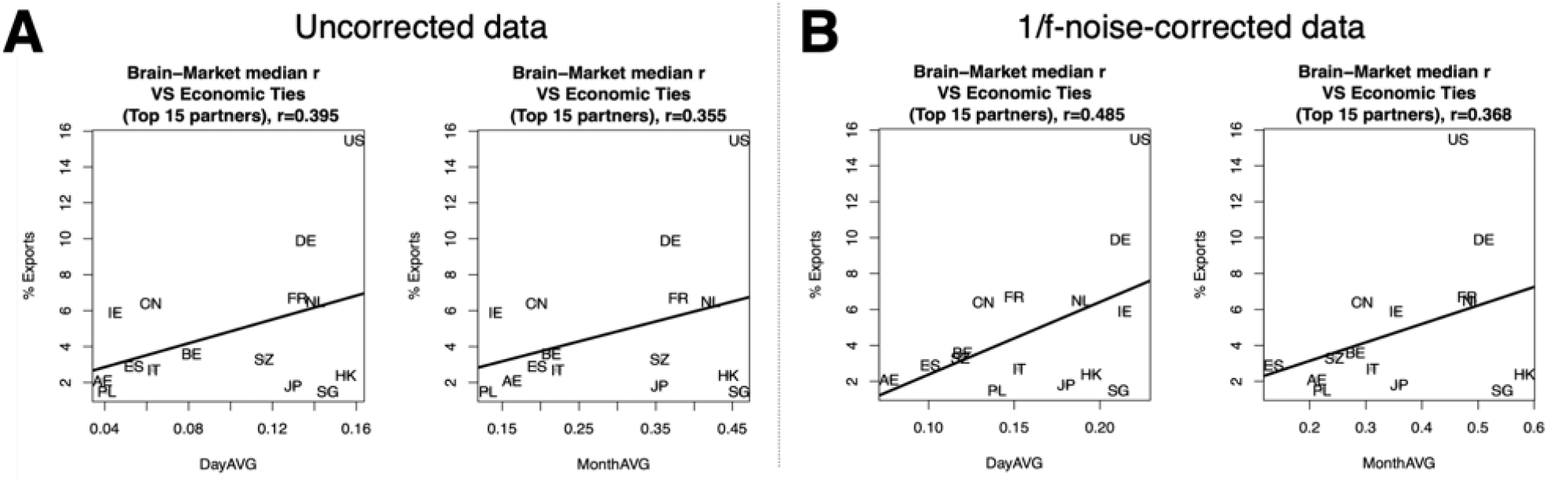
Association between Brain-Market link and economic ties with 15 UK’s top trading partners. The brain-market link measured as median Pearson correlation (r) for all of the 12 regions that exhibited significant associations in the main analysis was matched with markets of 15 UK’s top trading partners. The strength of economic ties was measured as a relative percent of all exports accessed from www.worldstopexports.com. Here we report the associations for the raw brain-market links. **A**), as well as the results following correction for 1/f noise. **B**), which entailed 1) simulation of the brain data with pink noise and 2) subtraction of the yielded correlations from the real data prior to calculating the medians. As expected, this procedure slightly increased the effect-sizes by increasing signal-to-noise ratio. Country labels: BE – Belgium, CN – China, FR – France, DE – Germany, HK – Hong Kong, IE – Ireland, IT – Italy, JP – Japan, NL – Netherlands, PL – Poland, SG – Singapore, ES – Spain, AE – United Arab Emirates, US – United States.

**Figure S7.**
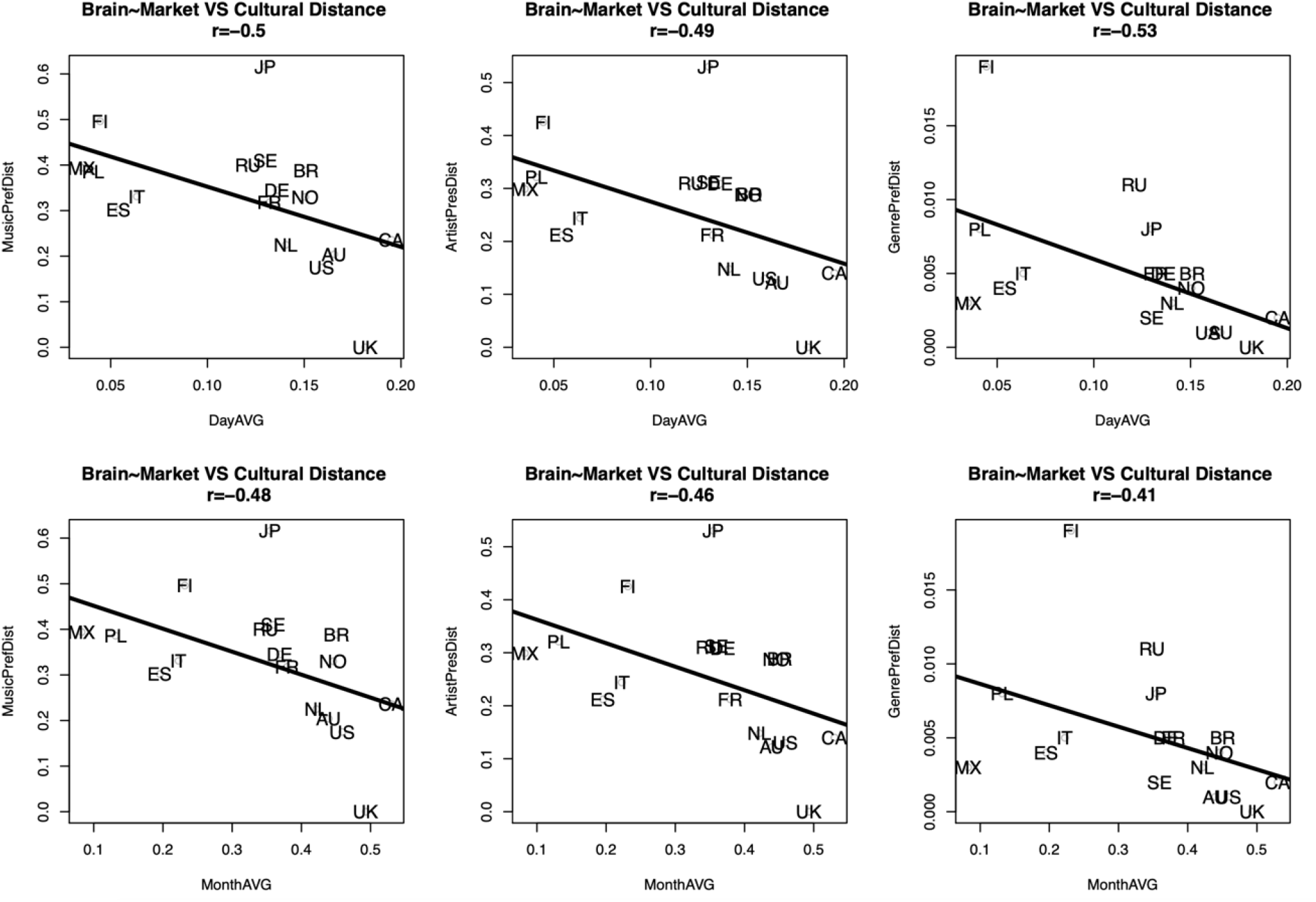
Association between Brain-Market link and sociocultural distances of the UK from 17 countries using data from Liu et al, 2018 ^43^. The brain-market link was measured as median Pearson correlation (r) for all of the 12 regions that exhibited significant associations in the main analysis. The sociocultural distance was calculated by Liu et al based on music (MusicPrefDist), artist (ArtistPrefDist) and genre preferences (MusicPrefDist). A negative relationship was found for the Brain-Market link and the afore-mentioned distances, i.e. the stronger the link the shorter the distance. The original study was focused on the following 20 countries: the United States (US), Russia (RU), Germany (DE), the United Kingdom (UK), Poland (PL), Brazil (BR), Finland (FI), Netherlands (NL), Spain (ES), Sweden (SE), Ukraine (market data not available), Canada (CA), France (FR), Australia (AU), Italy (IT), Japan (JP), Norway (NO), Mexico (MX), Czech Republic (market data not available), and Belarus (market data not available). The genre labels are artist-based and correspond to those in Allmusic, a major online music repository.

**Figure S8.**
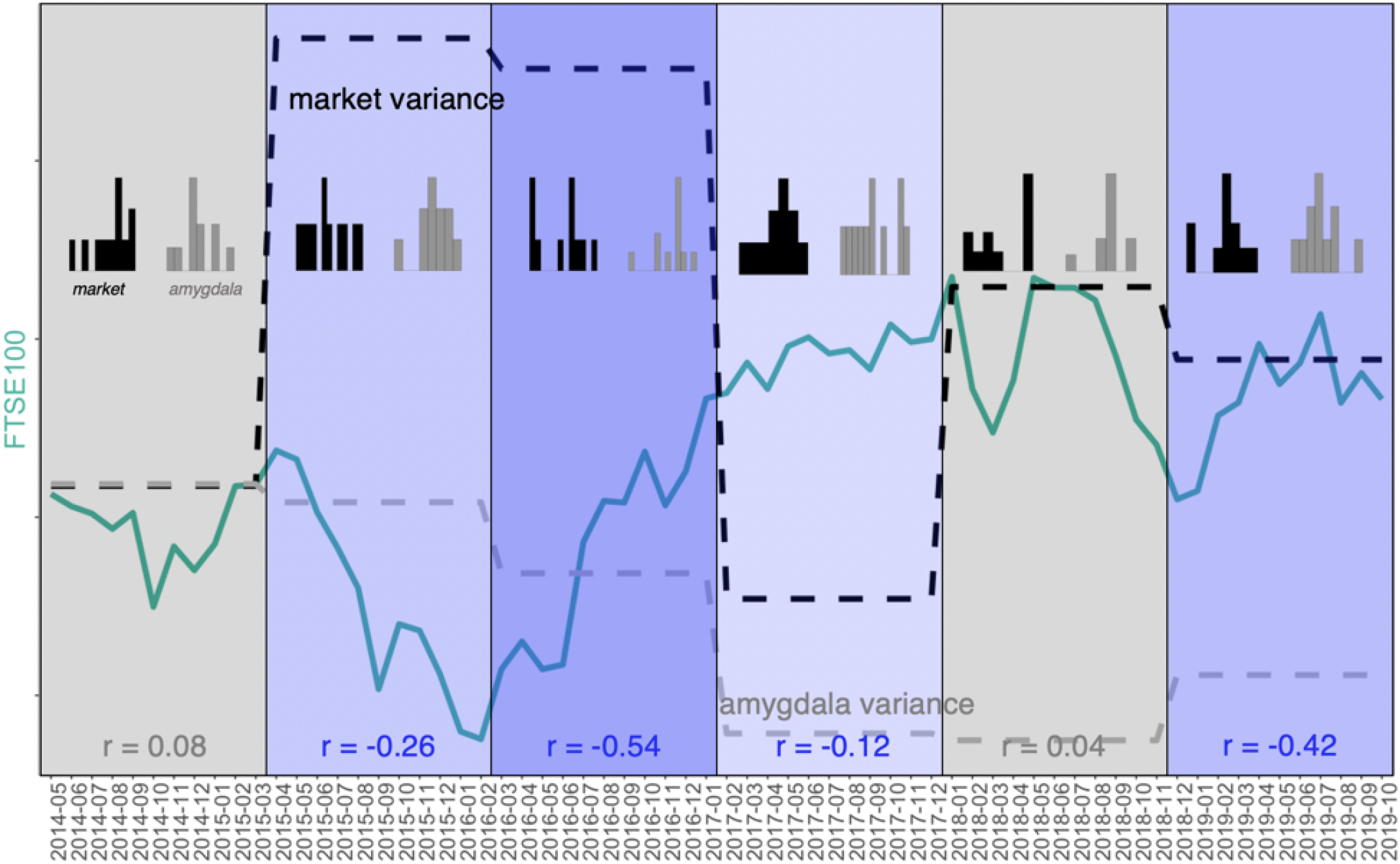
Correlation between amygdala volume and FTSE100 in different study periods. The timeline was split into 6 equal bins (11 months each) and correlations were calculated for each bin separately. The figure shows that the correlations are strongest during and following phase transition events, i.e. when the change and variability of stock market dynamics is most pronounced.

**Figure S9.**
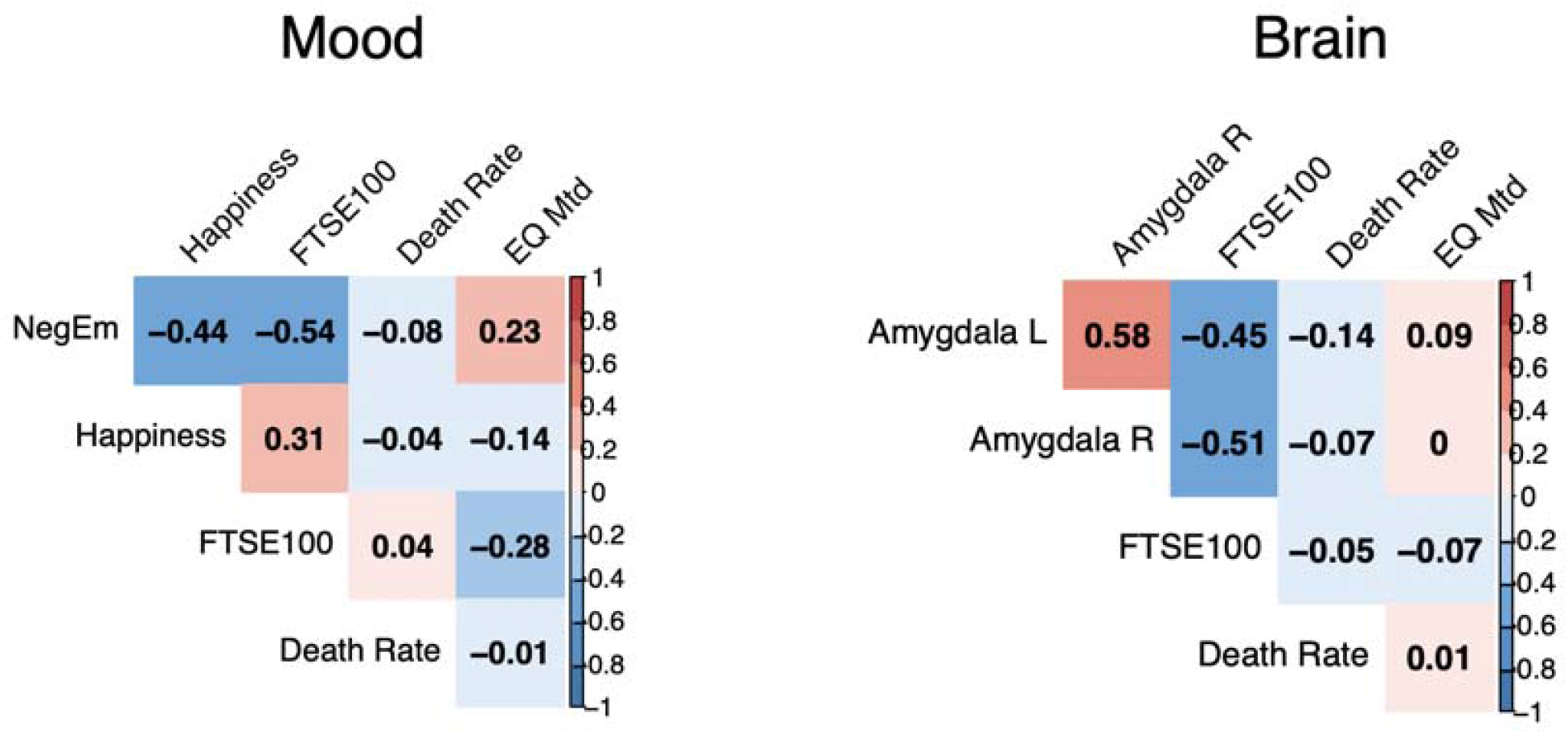
FTSE100 and alternative global candidate metrics with 1/f properties. The correlation plots show that compared to other global metrics FTSE100 exhibited strongest association with the investigated variables. All time-series were converted to a weekly scale for consistency (original scale of the public UK mortality data from www.mortality.org). Public UK seismic activity data measured on Richter magnitude scale (EQ Mtd) was accessed via earthquakes.bgs.ac.uk.

**Figure S10.**
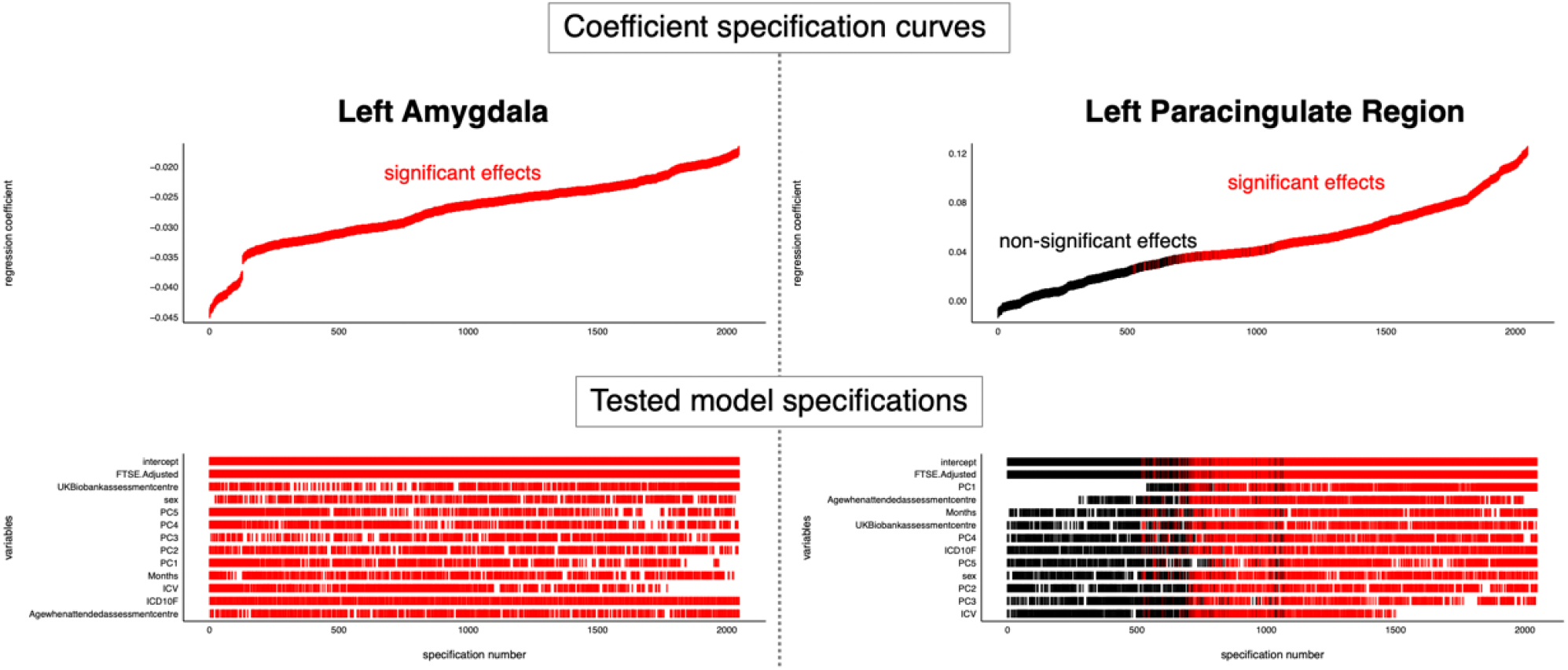
Specification curve analysis of the most and least significant brain regions from the main results. The analysis adhered to the Simonsohn’s protocol ^34^ focusing on the brain regions that had the strongest (amygdala) and weakest (paracingulate cortex) association with the FTSE100 index. The protocol entailed: 1) specification all reasonable models (to introduce all of the investigated nuisance covariates and their combinations), 2) plotting specification curves showing estimates/model fits as a function of analytic decisions, 3) testing how consistent the curve results are against a null hypothesis. The plot shows robustness of the investigated effects with respect to a wide variety of model specification strategies (i.e. most of the covariate combinations resulted in statistically significant estimates). All of the tested models fitted the data substantially better than a null model according to the AIC-criterion except for the one specifying non-adjusted effects of the FTSE100 on the paracingulate region, which, in line with the reported results, was equivalent to the null model (intercept only). All possible nested models were generated using the dredge() function from the MuMIn package.

**Figure S11.**
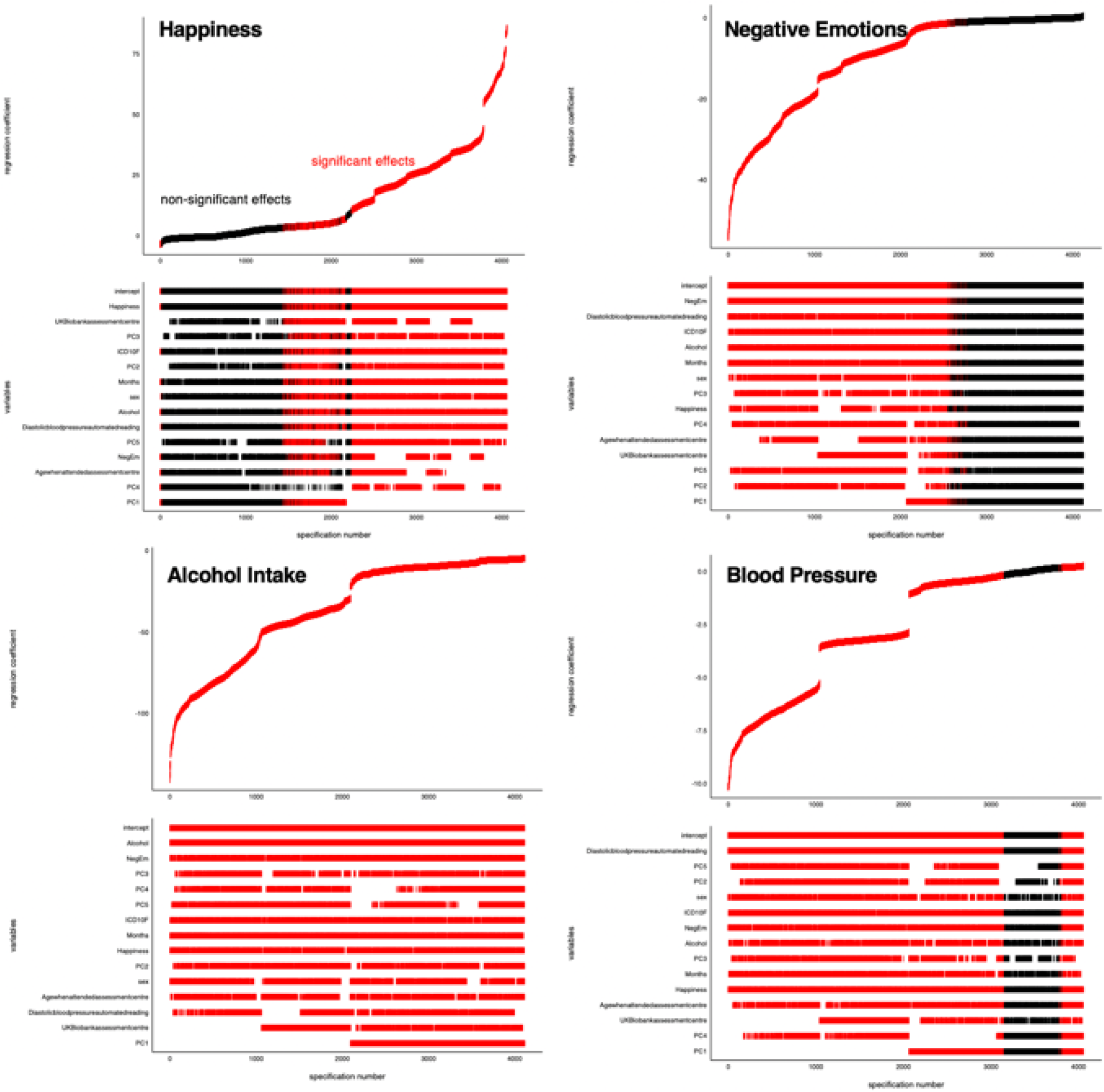
Specification curve analysis of the associations between FTSE100 and all of the investigated non-MRI variables: 14 years. For the present analysis FTSE100 was specified as a dependent variable in order to investigate independent variance contribution of all of the investigated non-MRI variables-of-interest (happiness, negative emotions, alcohol intake and diastolic blood pressure) and a number of confounders (non-UK stock markets, age, sex, psychiatric diagnosis, assessment center, seasonal effects) for the extended (14-year) period of the study. The plot shows stability of the investigated effects. All of the tested models fitted the data substantially better than a null model according to the AIC-criterion. A random 50% sample of all possible nested models were generated using the dredge() function from the MuMIn package.

**Figure S12.**
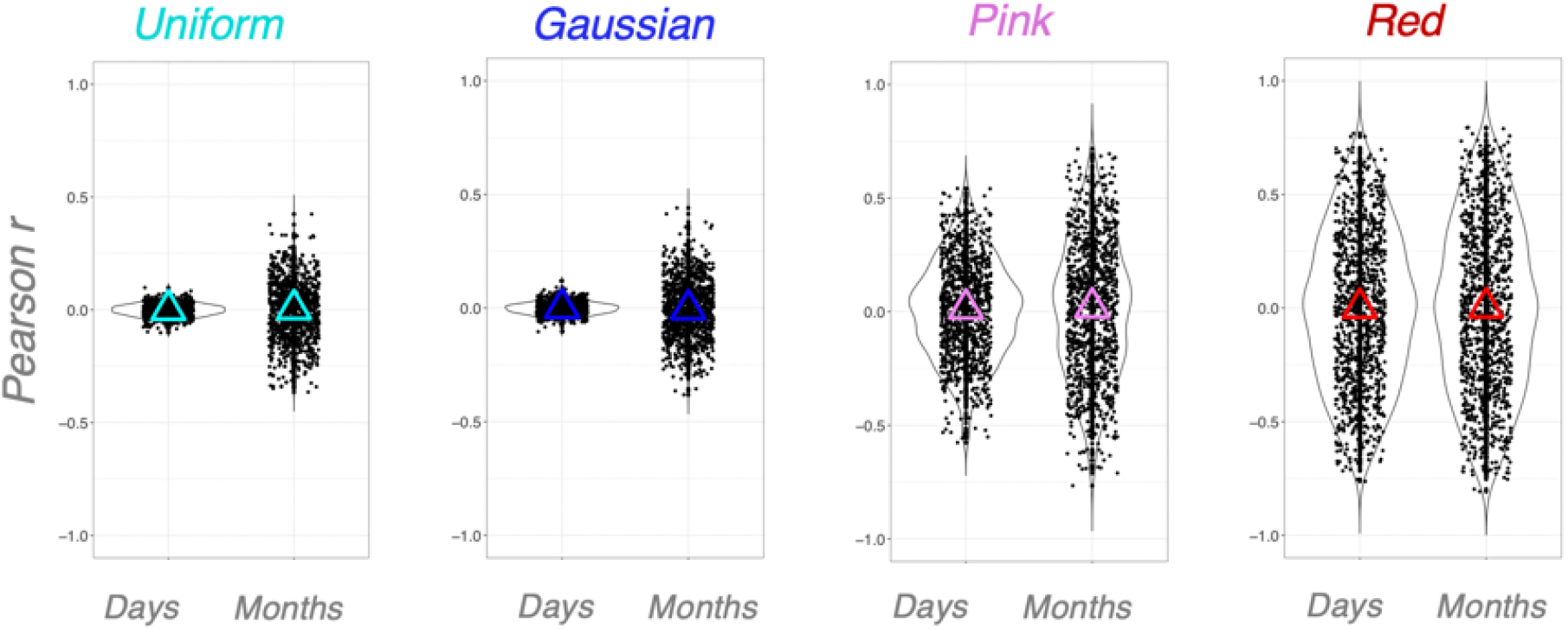
Raw noise simulation experiments. The plot shows convergence of the estimated effect-sizes to zero when no rootsquared-transformation is applied.

**Figure S13.**
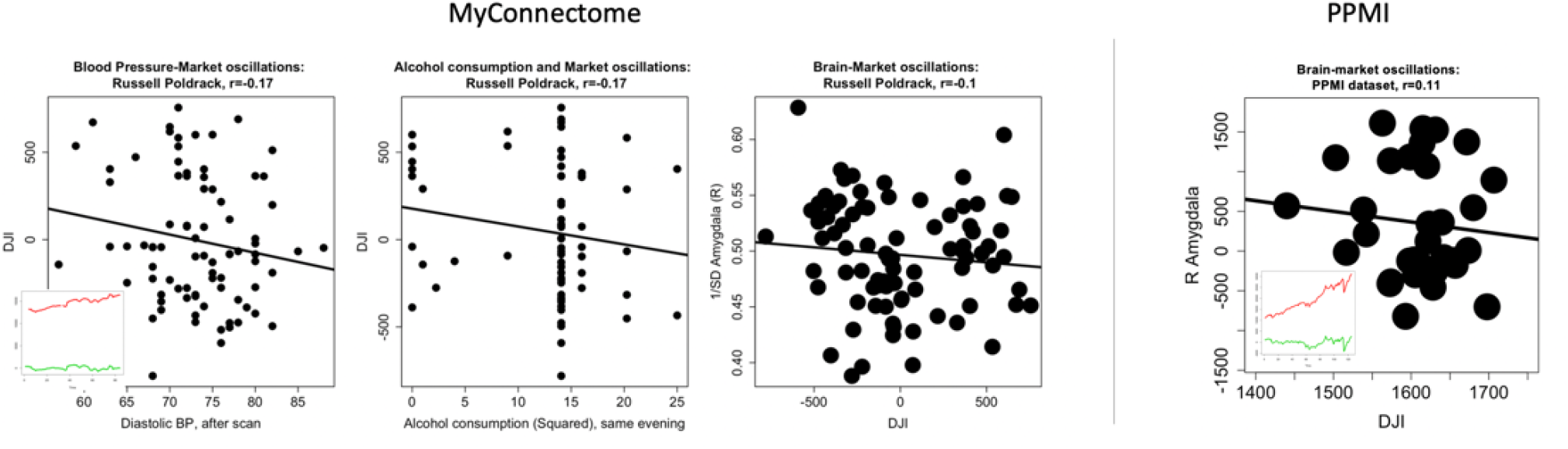
Detrended results in the MyConnectome and PPMI datasets. The plots demonstrate substantial reduction of effect-sizes with sustained directionality of the associations.

**Table S1.** **Study samples: descriptive statistics**

|  | Mood | Brain (main) |
| --- | --- | --- |
| Period, years (span) | 2006-03-13 - 2020-03-13 | 2014-05-02 - 2019-10-31 |
| Unique subjects / N, total | 479,791/ 547,005 | 39,755/41,182 |
| Age, years | 57.39(±8.35) | 63.67(±7.54) |
| Sex, Males/Females, %Males | 251635/295370 \| 46% M | 19434/21748 \| 47.19% M |
| Education, years | 16.43(±3.31) | 16.97(±2.63) |
| Intelligence score* | 6.17(±2.14) | 6.63(±2.06) |

\* Unweighted sum of the number of correct answers given to the 13 fluid intelligence questions.

**Table S2.**
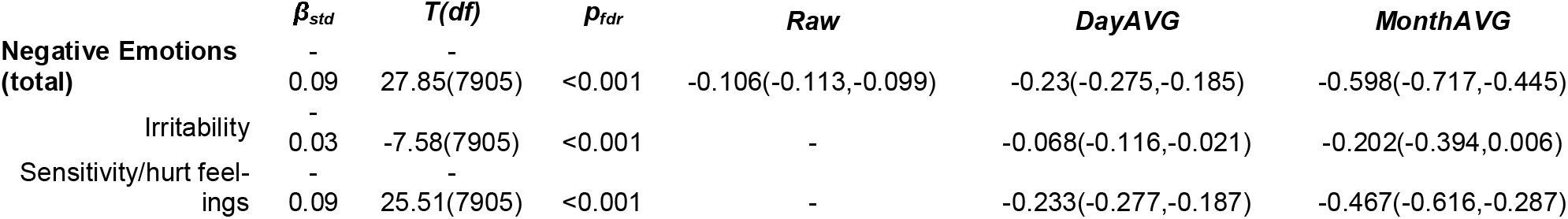
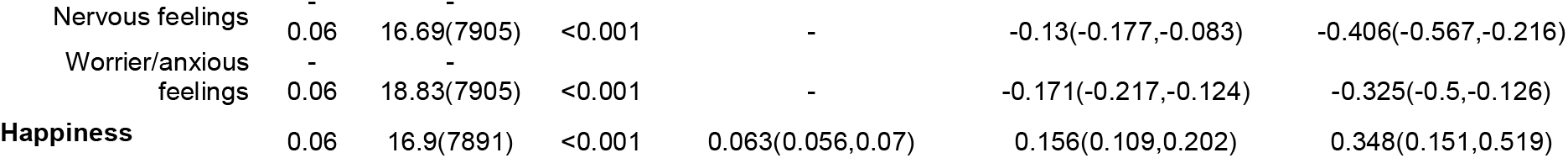
**Subjective well-being and FTSE100 scores: 5.5-year period.** The table shows stability of the market-mood relationships for the 5.5-year period (the same as the one investigated in the main analysis of brain-market relationships); β_std_ - standardized β coefficients, p_fdr_ – p-values corrected for multiple testing with false discovery rate. Subcomponents of negative emotions are binary variables (-), Day/MonthAVG – data averaged by days and months.

**Table S3.** **Main results adjusted for intracranial volume, demographics, psychiatric diagnosis and seasonal effects.** The table shows stability of the identified relationships when controlling for age, sex, psychiatric diagnosis, seasonal effects (months) and intracranial volume (cerebrospinal fluid, white and grey matter); ns – non-significant.

| Region | $\beta_{std}$ | $T_{776}$ | $p_{fdr}$ |
| --- | --- | --- | --- |
| **L Amygdala** | -0.049 | -6.24 | <0.001 |
| **R Amygdala** | -0.046 | -5.76 | <0.001 |
| **L Accumbens** | -0.021 | -2.92 | 0.008 |
| **R Accumbens** | -0.025 | -3.44 | 0.002 |
| **L LOFC** | -0.022 | -3.89 | 0.001 |
| **R LOFC** | -0.009 | -1.53 | ns |
| **L Insula** | 0.016 | 2.9 | 0.008 |
| **R Insula** | 0.02 | 3.47 | 0.002 |
| **L Subcallosal** | 0.013 | 2.29 | 0.042 |
| **R Subcallosal** | 0.011 | 1.82 | ns |
| **L Anterior Cingulate** | -0.002 | -0.26 | ns |
| **R Anterior Cingulate** | 0.005 | 0.78 | ns |
| **L Paracingulate** | 0.01 | 1.63 | ns |
| **R Paracingulate** | 0.011 | 1.78 | ns |

**Table S4.** **Main effects PCA-adjusted for the rest of the studied stock markets (first 5 principal components).** The table shows stability of the identified relationships when controlling for stock markets of the UK’s 15 top trading partners ^35^, measured as first 5 principal components extracted from the merged time-series.

| Region | Linear Mixed-Effects |  |  | Effect-sizes, Pearson r (95% C.I.) |  |  |
| --- | --- | --- | --- | --- | --- | --- |
| | $\beta_{std}$ | $T(df)$ | $p_{idr}$ | Raw, n=30,775 | DayAVG, n=1299 | MonthAVG, n=66 |
| *L Amygdala* | -0.056 | -4.2(878) | <0.001 | -0.01(-0.021.0.002) | -0.052(-0.106.0.002) | -0.131(-0.362.0.114) |
| *R Amygdala* | -0.061 | -4.96(878) | <0.001 | -0.012(-0.023.-0.001) | -0.066(-0.12.-0.012) | -0.206(-0.427.0.038) |
| *L Accumbens* | -0.014 | -1.1(878) | 0.311 | -0.001(-0.013.0.01) | -0.002(-0.057.0.052) | -0.029(-0.269.0.214) |
| *R Accumbens* | -0.01 | -0.75(878) | 0.488 | -0.002(-0.013.0.009) | -0.004(-0.058.0.051) | -0.052(-0.29.0.192) |
| *L LOFC* | 0.004 | 0.37(878) | 0.713 | <0.001(-0.011.0.011) | 0.005(-0.049.0.059) | 0.01(-0.233.0.251) |
| *R LOFC* | 0.022 | 1.81(878) | 0.088 | 0.003(-0.008.0.014) | 0.037(-0.018.0.091) | 0.101(-0.144.0.335) |
| *L Insula* | 0.069 | 5.64(878) | <0.001 | 0.01(-0.001.0.021) | 0.078(0.024.0.132) | 0.197(-0.047.0.419) |
| *R Insula* | 0.077 | 6.28(878) | <0.001 | 0.013(0.001.0.024) | 0.089(0.034.0.142) | 0.195(-0.049.0.417) |
| *L Subcallosal* | 0.056 | 4.52(878) | <0.001 | 0.01(-0.001.0.021) | 0.084(0.03.0.138) | 0.203(-0.041.0.424) |
| *R Subcallosal* | 0.048 | 3.88(878) | <0.001 | 0.009(-0.002.0.02) | 0.083(0.029.0.137) | 0.191(-0.053.0.414) |
| *L Anterior Cingulate* | 0.044 | 3.79(878) | <0.001 | 0.009(-0.002.0.02) | 0.071(0.017.0.125) | 0.133(-0.112.0.364) |
| *R Anterior Cingulate* | 0.047 | 3.93(878) | <0.001 | 0.009(-0.002.0.02) | 0.066(0.012.0.12) | 0.153(-0.092.0.381) |
| *L Paracingulate* | 0.047 | 3.92(878) | <0.001 | 0.009(-0.002.0.021) | 0.076(0.021.0.13) | 0.211(-0.033.0.431) |
| *R Paracingulate* | 0.041 | 3.4(878) | 0.001 | 0.008(-0.003.0.019) | 0.067(0.013.0.121) | 0.176(-0.069.0.401) |

**Table S5.** **Market-by-income interaction effect on the investigated volumetric brain measures, as well as the main of income.**

|  | Interaction (linear term) |  |  | Main Effect |  |  |
| --- | --- | --- | --- | --- | --- | --- |
| | $\beta$ | $T(df)$ | $p_{Fdr}$ | $\beta$ | $T(df)$ | $p_{Fdr}$ |
| **L Amygdala** | 0.01 | 1.24(776) | 0.516 | 0.03 | 4.39(776) | <0.001 |
| **R Amygdala** | <0.001 | 0.65(776) | 0.887 | 0.01 | 1.9(776) | 0.06 |
| **L Accumbens** | <0.001 | -0.81(776) | 0.832 | 0.08 | 14.32(776) | <0.001 |
| **R Accumbens** | <0.001 | -0.16(776) | 0.951 | 0.09 | 15.9(776) | <0.001 |
| **L LOFC** | -0.01 | -1.61(776) | 0.429 | 0.06 | 12.35(776) | <0.001 |
| **R LOFC\*** | -0.01 | -2.87(776) | 0.05 | 0.06 | 11.92(776) | <0.001 |
| **L Insula** | <0.001 | -0.25(776) | 0.951 | 0.02 | 4.32(776) | <0.001 |
| **R Insula** | <0.001 | 0.4(776) | 0.951 | 0.02 | 3.73(776) | <0.001 |
| **L Subcallosal** | -0.01 | -2.11(776) | 0.211 | 0.02 | 4.39(776) | <0.001 |
| **R Subcallosal** | -0.01 | -1.28(776) | 0.516 | <0.001 | -0.91(776) | 0.363 |
| **L Anterior Cingulate** | <0.001 | 0.33(776) | 0.951 | -0.03 | -5.88(776) | <0.001 |
| **R Anterior Cingulate** | <0.001 | -0.06(776) | 0.951 | -0.03 | -5.55(776) | <0.001 |
| **ICV** | -0.01 | -1.1(777) | 0.273 | 0.15 | 28.52(777) | <0.001 |

\*Post-hoc correlation tests revealed the largest effect-sizes in lowest and highest-income citizens

| **Income (n)** | **Pearson r (95% C.I.)** | **P-value** |
| --- | --- | --- |
| **Less than 18,000£ (4,557)** | -0.045(-0.078,-0.012) | 0.008 |
| **18,000-39,999£ (10,061)** | -0.018(-0.04,0.005) | 0.124 |
| **39,000-51,999£ (11,219)** | -0.033(-0.055,-0.012) | 0.002 |
| **52,000-100,000£ (8,570)** | -0.027(-0.052,-0.002) | 0.037 |
| **Greater than 100,000£ (2,699)** | -0.067(-0.11,-0.024) | 0.003 |

**Table S6.** **Brain-market associations: FTSE100 time-series deconvolved with Fast Fourier Transform (FFT) into low- and high-frequency bands.** Effects of low and high FTSE100 frequencies were estimated in one linear mixed-effects model assessing their independent contributions. The table shows that the main results are primarily driven by low-frequency oscillations, but high-frequency bands also exhibit some independent contribution.

|  | Low Frequency |  | High Frequency |  |
| --- | --- | --- | --- | --- |
| | $T_{881}$ | $P_{uncor}$ | $T_{881}$ | $P_{uncor}$ |
| **L Amygdala** | -9.59 | <0.001 | -0.63 | ns |
| **R Amygdala** | -11.71 | <0.001 | 0.91 | ns |
| **L Accumbens** | -9.78 | <0.001 | -0.44 | ns |
| **R Accumbens** | -11.25 | <0.001 | 0.33 | ns |
| **L LOFC** | -5.06 | <0.001 | -2.02 | 0.043 |
| **R LOFC** | -3.06 | 0.002 | -2.12 | 0.034 |
| **L Insula** | 11.96 | <0.001 | -1.46 | ns |
| **R Insula** | 10.62 | <0.001 | -1.86 | 0.06 |
| **L Subcallosal** | 7.01 | <0.001 | -2.25 | 0.02 |
| **R Subcallosal** | 6.16 | <0.001 | -1.89 | 0.059 |
| **L Anterior Cingulate** | 6.27 | <0.001 | -0.19 | ns |
| **R Anterior Cingulate** | 6.23 | <0.001 | -0.44 | ns |
| **L Paracingulate** | 1.91 | 0.056 | -0.93 | ns |
| **R Paracingulate** | 2.64 | 0.008 | -2.17 | 0.034 |

**Table S7.** **Autocorrelations in the investigated time-series** The Durbin-Watson (D-W) test revealed significant autocorrelations (ACs) present in the day-averaged population brain and market data. The ACs were detected in both time-series but were particularly strong in the stock market data (Hurst component for the FTSE100 data was estimated at 0.87, suggesting presence of long-term positive autocorrelation).

| Lag | LM: Brain~Market |  |  | LM: Market~Brain |  |  |
| --- | --- | --- | --- | --- | --- | --- |
|  | AC | D-W | p | AC | D-W | p |
| 1 | 0.05 | 1.898 | 0.004 | 0.851 | 0.295 | <0.001 |
| 2 | 0.002 | 1.994 | 0.82 | 0.842 | 0.313 | <0.001 |
| 3 | 0.014 | 1.966 | 0.344 | 0.841 | 0.312 | <0.001 |
| 4 | 0.017 | 1.958 | 0.296 | 0.84 | 0.314 | <0.001 |
| 5 | 0.022 | 1.948 | 0.162 | 0.837 | 0.318 | <0.001 |
| 6 | 0.029 | 1.932 | 0.062 | 0.837 | 0.318 | <0.001 |
| 7 | 0.054 | 1.882 | 0.002 | 0.839 | 0.313 | <0.001 |
| 8 | 0.095 | 1.8 | <0.001 | 0.844 | 0.303 | <0.001 |
| 9 | 0.004 | 1.981 | 0.636 | 0.829 | 0.334 | <0.001 |
| 10 | <0.001 | 1.989 | 0.908 | 0.827 | 0.337 | <0.001 |

**Table S8.** **Causal relationships between the studied brain variables and market oscillations (daily scale).** For all the regions that passed the Portmanteau stability test for residual serial correlation (highlighted in bold), hypothesis 2 (H2: “Market impacts Polulation Brain”) received slightly more support compared to hypothesis 1 (H2: “Population Brain impacts Market”) with the optimal lag length (L) determined according to AIC criterion. However, the H2 could not be ruled out, as it was also equivalently supported for amygdala and subcallosal cortex.

|  | **H1:**<br>"Population Brain impacts<br>Market" | **H2:**<br>"Market impacts Population Brain" | **Portmanteau**<br>**Stability Test** |
| --- | --- | --- | --- |
| **L Amygdala** | <u>$\chi^2(df)=21.72(5)$, $p=0.001$</u> | <u>$\chi^2(df)=26.28(5)$, $p<0.001$</u> | $L=5$, $\chi^2(df)=50.44(44)$, $p=0.234$ |
| **R Amygdala** | <u>$\chi^2(df)=26.5(13)$, $p=0.015$</u> | <u>$\chi^2(df)=35.76(13)$, $p=0.001$</u> | $L=13$, $\chi^2(df)=13.98(12)$, $p=0.302$ |
| **L Accumbens** | $\chi^2(df)=0.43(1)$, $p=0.512$ | $\chi^2(df)=2.49(1)$, $p=0.115$ | $L=1$, $\chi^2(df)=63.65(60)$, $p=0.349$ |
| **R Accumbens** | $\chi^2(df)=0.13(1)$, $p=0.718$ | <u>$\chi^2(df)=5.75(1)$, $p=0.016$</u> | $L=1$, $\chi^2(df)=72.39(60)$, $p=0.131$ |
| **L LOFC** | $\chi^2(df)=0.5(1)$, $p=0.481$ | $\chi^2(df)=2.92(1)$, $p=0.087$ | $L=1$, $\chi^2(df)=60.35(60)$, $p=0.463$ |
| **R LOFC** | $\chi^2(df)=0(1)$, $p=0.996$ | $\chi^2(df)=0.04(1)$, $p=0.841$ | $L=1$, $\chi^2(df)=65.54(60)$, $p=0.291$ |
| **L Insula** | $\chi^2(df)=13.57(11)$, $p=0.258$ | $\chi^2(df)=16.52(11)$, $p=0.123$ | $L=11$, $\chi^2(df)=20.86(20)$, $p=0.406$ |
| **R Insula** | $\chi^2(df)=13.65(11)$, $p=0.253$ | $\chi^2(df)=15.38(11)$, $p=0.166$ | $L=11$, $\chi^2(df)=25.92(20)$, $p=0.168$ |
| **L Subcallosal** | <u>$\chi^2(df)=18.82(8)$, $p=0.016$</u> | <u>$\chi^2(df)=16.31(8)$, $p=0.038$</u> | $L=8$, $\chi^2(df)=36.65(32)$, $p=0.262$ |
| **R Subcallosal** | <u>$\chi^2(df)=16.54(8)$, $p=0.035$</u> | <u>$\chi^2(df)=19.44(8)$, $p=0.013$</u> | $L=8$, $\chi^2(df)=37.55(32)$, $p=0.23$ |
| **L Anterior Cingulate** | $\chi^2(df)=7.19(5)$, $p=0.207$ | $\chi^2(df)=20.44(5)$, $p=0.001$ | $L=5$, $\chi^2(df)=61.26(44)$, $p=0.043$ |
| **R Anterior Cingulate** | $\chi^2(df)=0.05(1)$, $p=0.829$ | $\chi^2(df)=21.41(1)$, $p<0.001$ | $L=1$, $\chi^2(df)=116.67(60)$, $p<0.001$ |

**Table S9.** **Causal relationships between the studied mood variables and market oscillations (daily scale, 14 years).** For all the regions that passed the Portmanteau stability test for residual serial correlation (highlighted in bold), hypothesis 2 (H2: “Market impacts Polulation Mood”) received consistently more support compared to hypothesis 1 (H2: “Population Mood impacts Market”) with the optimal lag length (L) determined according to AIC criterion. Measures that passed the Portmanteau stability test for residual serial correlation are highlighted in bold. L - optimal lag length determined according to AIC criterion.

|  | **H1:**<br>**"Population Mood im-**<br>**pacts Market"** | **H2:**<br>**"Market impacts Popula-**<br>**tion Mood"** | **Portmanteau**<br>**Stability Test** |
| --- | --- | --- | --- |
| **NegEm** | $\chi^2(df)=2.38(2)$, $p=0.304$ | <u>$\chi^2(df)=7.1(2)$, $p=0.029$</u> | $L=2$, $\chi^2(df)=61.64(56)$, $p=0.281$ |
| **Irritability** | $\chi^2(df)=3.21(3)$, $p=0.36$ | <u>$\chi^2(df)=9.59(3)$, $p=0.022$</u> | $L=3$, $\chi^2(df)=63.43(52)$, $p=0.133$ |
| **Sensitivity/hurt feelings** | $\chi^2(df)=1.36(1)$, $p=0.243$ | <u>$\chi^2(df)=10.73(1)$, $p=0.001$</u> | $L=1$, $\chi^2(df)=69.56(60)$, $p=0.187$ |
| Nervous feelings | $\chi^2(df)=4.03(2)$, $p=0.133$ | $\chi^2(df)=0.16(2)$, $p=0.925$ | $L=2$, $\chi^2(df)=79.12(56)$, $p=0.023$ |
| Worrier/anxious feelings | $\chi^2(df)=0.06(1)$, $p=0.803$ | <u>$\chi^2(df)=10.4(1)$, $p=0.001$</u> | $L=1$, $\chi^2(df)=86.02(60)$, $p=0.015$ |
| **Happiness** | $\chi^2(df)=4.22(2)$, $p=0.121$ | <u>$\chi^2(df)=15.31(2)$, $p<0.001$</u> | $L=2$, $\chi^2(df)=69.92(56)$, $p=0.1$ |

**Table S10.** **Effects of autocorrelations on the studied associations.** The table demonstrates effects of autocorrelations present in stock market time-series. The effects are not present in the shuffled data, but appear for the 1.5-years-shifted market time-series. The same effect (including directional relationships) can be induced with 1/f noise, but it disappears after adjusting the results for the rest of the studied non-UK markets (first 5 Principal Components). *Note that when reporting effect-sizes of 1/f noise, we focused on a single simulation. Estimates from multiple simulations ultimately converge to zero (**Supplement Fig. S12**) due to inconsistent directionality of the associations.

|  | Shuffled |  |  | Shifted |  |  | 1/f Noise* |  |  | 1/f Noise* (PCA-corrected*) |  |  |
| --- | --- | --- | --- | --- | --- | --- | --- | --- | --- | --- | --- | --- |
| | $\beta_{std}$ | *T* | *Pfdr* | $\beta_{std}$ | *T* | *Pfdr* | $\beta_{std}$ | *T* | *Pfdr* | $\beta_{std}$ | *T* | *Pfdr* |
| L Amygdala | 0.002 | 0.38(810) | 0.928 | -0.015 | -2.81(758) | 0.006 | -0.005 | -0.93(758) | 0.35 | 0.009 | 1.34(563) | 0.261 |
| R Amygdala | -0.004 | -0.71(810) | 0.919 | -0.012 | -2.28(758) | 0.026 | -0.007 | -1.2(758) | 0.266 | 0.003 | 0.45(563) | 0.7 |
| L Accumbens | -0.005 | -0.89(810) | 0.919 | -0.052 | -10.49(758) | <0.001 | -0.021 | -3.88(758) | <0.001 | -0.004 | -0.56(563) | 0.666 |
| R Accumbens | <0.001 | 0.09(810) | 0.928 | -0.053 | -10.67(758) | <0.001 | -0.026 | -4.75(758) | <0.001 | -0.009 | -1.42(563) | 0.261 |
| L LOFC | -0.002 | -0.51(810) | 0.919 | -0.028 | -8.37(758) | <0.001 | -0.018 | -3.82(758) | <0.001 | -0.007 | -1.31(563) | 0.261 |
| R LOFC | <0.001 | -0.1(810) | 0.928 | -0.036 | -10.74(758) | <0.001 | -0.023 | -4.84(758) | <0.001 | -0.008 | -1.51(563) | 0.249 |
| L Insula | -0.002 | -0.53(810) | 0.919 | -0.002 | -0.49(758) | 0.623 | -0.008 | -1.78(758) | 0.113 | -0.017 | -2.9(563) | 0.046 |
| R Insula | -0.001 | -0.15(810) | 0.928 | -0.006 | -1.63(758) | 0.111 | -0.008 | -1.68(758) | 0.118 | -0.014 | -2.51(563) | 0.046 |
| L Subcallosal | -0.003 | -0.54(810) | 0.919 | -0.022 | -6.04(758) | <0.001 | -0.016 | -3.35(758) | 0.002 | -0.015 | -2.61(563) | 0.046 |
| R Subcallosal | -0.003 | -0.53(810) | 0.919 | -0.012 | -3.3(758) | 0.002 | -0.01 | -2.04(758) | 0.069 | -0.014 | -2.39(563) | 0.051 |
| L Anterior Cingulate | 0.003 | 0.74(810) | 0.919 | 0.01 | 3.23(758) | 0.002 | 0.004 | 1.05(758) | 0.317 | -0.004 | -0.7(563) | 0.602 |
| R Anterior Cingulate | 0.005 | 1.07(810) | 0.919 | 0.009 | 2.81(758) | 0.006 | 0.007 | 1.68(758) | 0.118 | -0.001 | -0.25(563) | 0.805 |
| L Paracingulate | -0.001 | -0.24(810) | 0.928 | -0.031 | -9.81(758) | <0.001 | -0.018 | -4.13(758) | <0.001 | -0.009 | -1.7(563) | 0.225 |
| R Paracingulate | -0.002 | -0.51(810) | 0.919 | -0.031 | -9.61(758) | <0.001 | -0.021 | -4.72(758) | <0.001 | -0.015 | -2.75(563) | 0.046 |
| Intracranial Volume | -0.004 | -1.16(810) | 0.919 | -0.033 | -17.86(758) | <0.001 | -0.024 | -7.51(758) | <0.001 | -0.006 | -1.5(563) | 0.249 |

**Table S11.** **Associations between FTSE100 and the main investigated variables tested with mutual information criterion.** Two values per test are provided for the non-brain variables: 14 years (full dataset) / 5.5 years (MRI subsample). Subcomponents of negative emotions are binary variables (-), Day/MonthAVG – data averaged by days and months.

|  | Raw | DayAVG | MonthAVG |
| --- | --- | --- | --- |
| Negative Emotions (total) | 0.003/0.004 | 0.254/0.049 | 0.461/0.136 |
| Irritability | - | 0.172/0.059 | 0.23/0.178 |
| Sensitivity/hurt feelings | - | 0.262/0.081 | 0.522/0.255 |
| Nervous feelings | - | 0.216/0.078 | 0.415/0.066 |
| Worrier/anxious feelings | - | 0.224/0.047 | 0.459/0.197 |
| Happiness | 0.001/0.002 | 0.145/0.078 | 0.243/0.061 |
| Diastolic blood pressure | 0.007/0.014 | 0.23/0.049 | 0.363/0.134 |
| Alcohol (intake frequency) | 0.002/0.003 | 0.132/0.056 | 0.236/0.093 |
| Alcohol (composite score intake) | 0.008/0.015 | 0.167/0.064 | 0.36/0.058 |
| L Amygdala | 0.018 | 0.111 | 0.317 |
| R Amygdala | 0.018 | 0.119 | 0.314 |
| L Accumbens | 0.018 | 0.096 | 0.432 |
| R Accumbens | 0.019 | 0.106 | 0.556 |
| L LOFC | 0.017 | 0.085 | 0.191 |
| R LOFC | 0.016 | 0.067 | 0.173 |
| L Insula | 0.016 | 0.091 | 0.294 |
| *R Insula* | 0.016 | 0.078 | 0.209 |
| *L Subcallosal* | 0.015 | 0.083 | 0.138 |
| *R Subcallosal* | 0.014 | 0.072 | 0.138 |
| *L Anterior Cingulate* | 0.016 | 0.1 | 0.267 |
| *R Anterior Cingulate* | 0.016 | 0.091 | 0.256 |
| *L Paracingulate* | 0.016 | 0.066 | 0.051 |
| *R Paracingulate* | 0.017 | 0.066 | 0.143 |

## Acknowledgments

The authors would like to thank Enzo Tagliazucchi, Henrik Larsson, Ralf Kuja-Halkola, Otilia Horntvedt, Markus Hjorth, Madelene Holm, Andres Cabrera and Irina Letiagina for the valuable discussions and exchange of ideas. We also thank William Thompson for feedback on the analysis, code review and commenting on earlier drafts of the manuscript.

MRI data used to replicate the main findings were obtained from the Parkinson’s Progression Markers Initiative (PPMI) database (www.ppmi-info.org). PPMI Data and Publications Committee approved the manuscript before the submission.

## Author contributions

AL – formulated the main hypothesis, prepared the first draft of the UK Biobank data access application, preregistered the study, conducted analyses, interpreted results, produced initial draft of the manuscript; GD,MK – suggested specification curve analyses, contributed in results interpretations, discussions and drafting; CA – was closely involved in results interpretations and discussions, proposed a number of supplementary analyses, contributed in drafting; KA – replicated main results, contributed in creating figures, took part in discussions and results interpretations, contributed in manuscript drafting; MI – was closely involved in results interpretation and discussions, proposed several important analyses, drafted the manuscript. PP – acquired funding, was closely involved in hypotheses submissions and preparations of the UK Biobank application, preregistration, proposed a number of important secondary and supplementary analyses, played key roles in results interpretation and discussions. All authors intellectually contributed to the study and took active parts in drafting and manuscript preparations, and approved the final draft of the manuscript.

## Data and materials availability

The access to the UK Biobank data was granted to the authors after submitting project description with stated hypotheses and analysis plan. The study was preregistered at the Open Science Foundation Framework database (https://osf.io/h52gk) prior to data transfer.

UK Biobank remains the owner of the database and accepts data request from third parties after approving corresponding project proposals and payments of the data access fees (more details: https://www.ukbiobank.ac.uk/principles-of-access). Similarly, Parkinson’s Progression Markers Initiative MRI data used to replicate the main findings is not publicly open, but can be accessed after completing a corresponding registration form (https://www.ppmi-info.org). Main data analysis steps are illustrated with MyConnectome data in a reproducible R-script attached to the submission (“GETAB_Poldrack.R”).

